# Multiple energy pathways structure mercury biomagnification in an Amazonian River food web

**DOI:** 10.64898/2026.09.28.755185

**Authors:** William Gonzalez-Daza, Angélica María Torres-Bejarano, Juan Camilo Ríos-Orjuela, Santiago R. Duque

## Abstract

Mercury biomagnification in Amazonian rivers is usually assessed through fishes, yet contaminants can also move across riparian food webs through multiple energy pathways. We evaluated how total mercury (THg) is structured across the upper Caquetá River basin, a mining-impacted river–floodplain system in the Colombian Amazon. Using stable carbon and nitrogen isotopes, trophic guild classification, biomagnification models, and Bayesian mixing models, we integrated basal resources, macroinvertebrates, fishes, and birds to link mercury exposure with food-web architecture. THg increased with trophic enrichment in fishes, birds, and the combined food web, indicating biomagnification across aquatic and aquatic-associated consumers. Reconstructed pathways showed that phytoplankton and detritus support intermediate fish consumers, which in turn connect basal production to carnivore–piscivorous fishes and aquatic predatory birds. Avian results suggest that mercury exposure is not restricted to aquatic predators, as hummingbirds showed unexpectedly elevated THg values that may reflect additional riparian or arthropod-mediated pathways. Overall, mercury transfer in the upper Caquetá emerges from the interaction between trophic position and energy-flow structure, highlighting the need to evaluate contamination magnification through integrated aquatic–riparian food webs rather than single consumer groups.

## Introduction

Mercury (Hg) is a priority environmental pollutant because of its persistence, toxicity, and long-range transport (Marrugo-Negrete et al., 2008; Olivero-Verbel et al., 2016). Although released naturally through volcanic activity and rock weathering, anthropogenic activities—particularly gold mining, industry, and fossil-fuel combustion—have substantially increased environmental Hg inputs (Laffont et al., 2021; Olivero-Verbel et al., 2016; Pouilly et al., 2013). Once released, Hg can persist for decades in soils and sediments and be transported through river networks, spreading contamination beyond mining areas to remote ecosystems and aquatic-resource-dependent populations (Lacerda, 1997; Gasca Álvarez, 2000; Veiga & Hinton, 2002; Alcala-Orozco et al., 2020; Meneses et al., 2022).

Hg risk largely depends on microbial methylation into monomethylmercury (MeHg), a highly toxic and bioavailable form produced mainly by sulfate- and iron-reducing bacteria and methanogenic archaea (Gilmour et al., 2013; McDaniel et al., 2020; Cardona et al., 2022; Casso-Hartmann et al., 2022). Methylation is favored in tropical flooded soils, sediments, and periphyton-rich habitats, where low oxygen and abundant organic matter provide suitable conditions (Achá et al., 2011; Paranjape & Hall, 2017). Because MeHg is efficiently assimilated and retained in tissues, it readily bioaccumulates and biomagnifies through food webs (Coelho et al., 2013; Vargas Licona & Marrugo-Negrete, 2019).

Hg concentrations generally increase from primary producers to top predators, with piscivorous fish often exceeding recommended human-consumption thresholds (Lavoie et al., 2013; Mussy et al., 2023; Pouilly et al., 2013). Fish integrate exposure across time and trophic pathways and transfer Hg to wildlife and humans (Passos & Mergler, 2008; Selin, 2009). Accordingly, trophic position is a strong predictor of fish Hg concentrations in the Amazon basin, where carnivorous species usually exhibit the highest burdens (Barbosa et al., 2003; Olivero-Verbel et al., 2016; Alcala-Orozco et al., 2020; Crespo-Lopez et al., 2021).

However, trophic position alone may not explain Hg bioaccumulation. Basal energy pathways, particularly benthic versus pelagic carbon contributions, can also regulate contaminant transfer (Coelho et al., 2013; Passos & Mergler, 2008; Paiva et al., 2024). Because methylation is spatially heterogeneous and enhanced in sediments, flooded soils, and detritus-rich habitats (Ullrich et al., 2001; Chen et al., 2014), benthic- or detritus-supported food webs may expose consumers to more Hg than pelagic pathways (Riva-Murray et al., 2013; Rocha et al., 2015). This distinction is especially relevant in tropical rivers, where hydrological variability and allochthonous inputs generate complex mixtures of carbon sources (Laffont et al., 2021).

Stable carbon (δ¹³C) and nitrogen (δ¹ N) isotopes can determine whether Hg transfer reflects trophic position, basal energy pathways, or both. δ¹ N indicates trophic position through its enrichment across trophic levels (Post, 2002; Vander Zanden & Rasmussen, 2001), whereas δ¹³C helps distinguish terrestrial C3 inputs, aquatic macrophytes, periphyton, and phytoplankton (Peterson & Fry, 1987; Forsberg et al., 1993). Their integration with Hg measurements allows trophic magnification and contaminant pathways to be evaluated simultaneously (Lavoie et al., 2013). Moreover, δ¹³C can explain Hg variation independently of trophic position, highlighting the importance of basal-resource structure (Riva-Murray et al., 2013; Swinton et al., 2022).

Despite these advances, tropical Hg studies have focused mainly on fish and rarely include upper-level consumers such as birds. It therefore remains unclear whether aquatic biomagnification patterns extend to predators that integrate diverse prey across broader spatial and temporal scales. Aquatic and semi-aquatic birds occupy high trophic positions, accumulate substantial Hg burdens, and serve as sentinels of local contamination and broader ecological processes (Evers et al., 2005; Clayden et al., 2015; Sayers et al., 2023). In Neotropical mining regions, their Hg concentrations may reach toxicologically concerning levels, particularly in piscivorous and insectivorous species (Sierra-Marquez et al., 2018; Pisconte et al., 2024).

In the Amazon, artisanal and small-scale gold mining (ASGM) is the principal anthropogenic Hg source (Alcala-Orozco et al., 2019; Martoredjo et al., 2024). The upper Caquetá River basin in Colombia is particularly relevant because ASGM and land-use change have intensified, with Hg contamination documented in aquatic ecosystems and human populations (Olivero-Verbel et al., 2016; Araruna et al., 2025). Its diverse food webs integrate aquatic and terrestrial energy sources, generating substantial heterogeneity in carbon use and contaminant exposure.

Here, we evaluate total mercury (THg) variation across the upper Caquetá River food web and examine how isotopically inferred energy pathways connect aquatic Hg exposure to fish and birds. By integrating THg, stable isotopes, trophic guilds, and Bayesian mixing models, we test whether THg increases with δ¹ N-inferred trophic enrichment and identify the pathways through which Hg reaches upper-level consumers.

We hypothesize that THg increases with δ¹ N and trophic position, indicating biomagnification across the food web (H1). We further predict that δ¹³C- and δ¹ N-based mixing models will reveal distinct pathways connecting basal resources, fish guilds, and aquatic or riparian birds, demonstrating that Hg transfer also depends on basal-resource use (H2). Finally, we expect piscivorous fish and aquatic birds to exhibit the highest THg concentrations and act as effective sentinels of ecosystem contamination (H3).

## Materials and Methods

### Study area

The study was conducted in the upper Caquetá River basin, Colombian Amazon, at Curillo and San Antonio de Getuchá (Caquetá Department) and Mecaya (Putumayo Department). Sampling was conducted during the rainy season (June 2024) in rivers, streams, and floodplain lakes associated with the Caquetá and Ortegüaza river systems (Figure 1).

**Figure 1.**
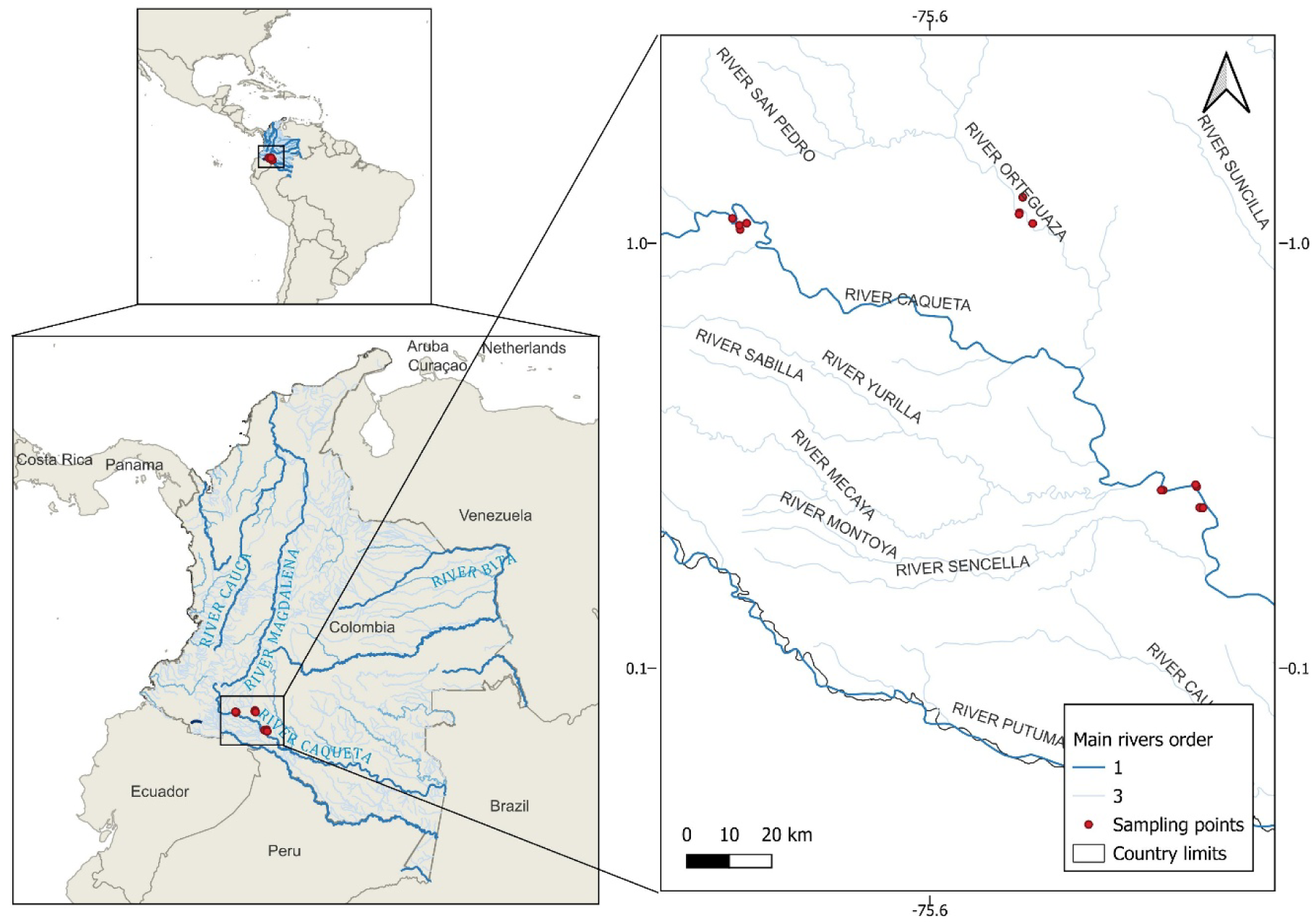
Location of the study area and sampling sites (red dots) in the Caquetá River basin, Colombian Amazon. Major rivers, stream order, and international boundaries are shown.

The region has a humid tropical climate, with annual precipitation exceeding 3,300 mm, mean temperatures of approximately 25°C, and relative humidity above 80%. The Caquetá is a slightly acidic, weakly mineralized whitewater river with high dissolved oxygen concentrations. Artisanal alluvial gold mining has occurred throughout the basin for approximately two decades and represents an important source of Hg contamination.

### Sampling design

Sediments, basal resources, macroinvertebrates, fishes, and birds were sampled to characterize trophic pathways and Hg transfer across aquatic and riparian food webs. Sediments were collected with an Ekman dredge, refrigerated, and analyzed for total mercury (THg). Periphyton was scraped from natural substrates, phytoplankton was collected using horizontal net tows (24-μm mesh), and aquatic macrophytes were obtained from dominant vegetation patches. Macroinvertebrates associated with macrophytes were sampled using D-frame nets and identified to the lowest feasible taxonomic level.

Fish sampling targeted locally consumed species potentially exposed to Hg bioaccumulation. Specimens were collected from lotic and lentic habitats using cast nets, gillnets, seine nets, baited traps, and longlines; additional specimens were obtained from local fishers. Fish were identified and measured, and muscle tissue was collected for THg and stable isotope analyses. Stomach contents were classified into major dietary categories.

Birds were captured using mist nets in primary forest, riparian vegetation, pasture–crop mosaics, and pasture mosaics with natural vegetation. Contour feathers were collected for THg and stable isotope analyses, after which individuals were released at the capture site.

### Trophic guild classification

Bird species were classified as invertivores, omnivores, frugi-granivores, nectarivores, or aquatic predators according to their dominant diet, foraging behavior, habitat use, and association with aquatic environments. Classifications were based on EltonTraits 1.0 (Wilman et al., 2014), Birds of the World (Billerman et al., 2026), and regional literature (Lopes et al., 2016). Stable isotope values were used only to assess consistency. The detailed protocol is provided in the Supplementary Material.

Fish trophic guilds were defined by integrating stomach-content composition, δ¹³C and δ¹ N values, total mercury (THg) concentrations, and standard length. Dietary items were grouped into fish, aquatic macroinvertebrates, detritus, aquatic or terrestrial plants, terrestrial insects, unidentified animal tissue, and empty stomachs. Mean dietary proportions were calculated for each species, while stable isotope signatures, THg concentrations, and body size provided complementary information on carbon sources, trophic position, contaminant accumulation, and ontogenetic variation. THg concentrations and standard length were log□□-transformed and standardized before analysis.

Bray-Curtis dissimilarities were calculated from species-level dietary proportions, whereas Euclidean distances were calculated from standardized δ¹³C, δ¹ N, log□□-transformed THg, and log□□-transformed standard length. These distance matrices were combined into a mixed trophic dissimilarity matrix as follows:

where D_diet_ represents dietary dissimilarity, D_traits_ represents dissimilarity based on isotopic and biological variables, and w_diet_ and w_traits_ are their respective weights. Fish species were subsequently grouped using average-linkage hierarchical clustering (UPGMA). Species connected at lower linkage heights were considered to have more similar feeding strategies, whereas those joined at higher linkage heights exhibited greater trophic differentiation. The resulting clusters were interpreted according to their dominant dietary resources and evaluated using isotopic position, THg concentrations, and body size. These clusters were then used to classify species into trophic guilds for Bayesian mixing models (MixSIAR) and mercury biomagnification analyses.

### Laboratory analyses

Total mercury (THg) concentrations in fish and feathers were determined using a direct mercury analyzer (RA-915 LAB, Lumex Instruments; Rumiantseva et al., 2022; Bazhenova et al., 2024), based on thermal decomposition and atomic absorption spectrometry with Zeeman background correction. Animal and plant tissues were freeze-dried and analyzed using 10-mg aliquots; sediments were air-dried for 24 h, pulverized, and analyzed using 200-mg aliquots, whereas feathers and hair were finely cut and analyzed using 20-mg aliquots. Water samples were analyzed directly using 100 μL aliquots. Analytical quality assurance and quality control followed EURACHEM, EPA, IDEAM, ONAC, and Standard Methods guidelines. Certified reference materials were used to assess analytical accuracy, including IAEA-436A for fish, IAEA-461 for macroinvertebrates, IAEA-450 for macrophytes, periphyton and detritus, IAEA-475 for sediments, IAEA-086 for human hair, and a High Purity Standards Hg solution for water. Recovery and repeatability were evaluated using five replicate measurements, with mean recovery, standard deviation, and coefficient of variation calculated for each matrix. Method detection limits ranged from 0.027 to 0.028 ng for biological matrices and were 0.20 ng for sediments, while quantification limits ranged from 0.083 to 0.086 ng and were 0.62 ng for sediments, respectively. Expanded measurement uncertainty ranged from 0.053 to 0.056 ng for biological matrices. THg concentrations were expressed as mg kg ¹ dry weight.

Stable isotope analyses of carbon (δ¹³C) and nitrogen (δ¹ N) were performed using an elemental analyzer coupled to an isotope ratio mass spectrometer (EA-IRMS) at the *Centro de Isótopos Estáveis Prof. Dr. Carlos Ducatti*, Instituto de Biociências, UNESP, Campus Botucatu, Brazil. Results were expressed in ‰ relative to Vienna Pee Dee Belemnite (V-PDB) for carbon and atmospheric N (AIR) for nitrogen. Analytical precision was ≤ 0.1‰ for δ¹³C and ≤ 0.2‰ for δ¹ N, verified through replicate analyses of internal standards calibrated against IAEA reference materials.

### Sample preparation and lipid correction

Fish muscle samples were dried, ground, and analyzed in duplicate. When muscle C:N ratios exceeded 3.5, δ¹³C values were corrected using the linear model of Post et al. (2007) (δ¹³C_corrected = δ¹³C_untreated − 3.32 + 0.99 × C:N) to account for lipid depletion in ¹³C. Feathers were cleaned by soaking in a 2:1 chloroform:methanol solution for 24 h, rinsed twice with the same mixture, and air-dried in a fume hood for at least 48 h before analysis.

### Mercury biomagnification analyses

Biomagnification was evaluated separately for fishes, birds, and the combined fish–bird dataset. Because THg concentrations were right-skewed, they were log□□-transformed and modeled as:

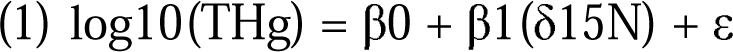

where δ¹ N represented trophic position and log□□(THg) represented mercury concentration.

Positive and significant slopes were interpreted as evidence of biomagnification. Model performance was assessed using coefficients of determination (R²), ANOVA, and significance tests.

### Bayesian stable isotope mixing models

Resource contributions to fish and bird guilds were estimated using Bayesian mixing models implemented in MixSIAR. Because locality-level sample sizes were limited, data from Curillo, Mecaya, and San Antonio de Getuchá were pooled to reconstruct regional trophic pathways. Mixing spaces were examined before model fitting to assess source separation, source–consumer relationships, and isotopic overlap. Candidate sources were retained based on isotopic distinction, stomach contents, field observations, and ecological relevance.

Fish were grouped as primary consumers, invertivores, omnivores, and carnivore–piscivores. Their respective sources were: (1) detritus, phytoplankton, macrophytes, and periphyton; (2) detritus, allochthonous insects, and aquatic macroinvertebrates; (3) detritus, phytoplankton, insects, macrophytes, and aquatic macroinvertebrates; and (4) primary-consumer, invertivorous, and omnivorous fishes.

Birds were grouped as invertivores, omnivores, frugi-granivores, nectarivores, and aquatic predators. Their sources were: (1) insects and aquatic macroinvertebrates; (2) insects, aquatic macroinvertebrates, and terrestrial plants; (3) terrestrial plants and insects; (4) terrestrial plants, insects, and aquatic macroinvertebrates; and (5) fish guilds and aquatic macroinvertebrates.

All Bayesian stable isotope mixing models were fitted in R version 4.5.3 using MixSIAR version 3.1.12 in RStudio 2024.04.2 (Build 764). Models incorporated both residual and process error structures and used trophic enrichment factors of 0.4 ± 0.2‰ for δ¹³C and 2.8 ± 0.5‰ for δ¹ N (mean ± SD). Posterior distributions were estimated through Markov Chain Monte Carlo simulations using uninformative priors and MixSIAR’s “normal” configuration, comprising three Markov chains of 100,000 iterations each, a burn-in of 50,000 iterations, and a thinning interval of 50. Model convergence was assessed using the Gelman–Rubin and Geweke diagnostics (Geweke, 1991; Gelman et al., 2014; Guerrero & Rogers, 2020). Posterior means, standard deviations, and 95% credible intervals were used to quantify the proportional contributions of resources and identify the dominant trophic pathways.

### Trophic network reconstruction

Posterior mean contributions from MixSIAR were used to construct directed trophic networks linking basal resources, intermediate consumers, fishes, and birds. Edges were weighted by posterior mean contribution, and only links contributing ≥10% were retained.

Nodes were arranged into four trophic levels: basal resources, primary consumers or prey, intermediate consumers, and top predators. Basal resources included detritus, phytoplankton, periphyton, macrophytes, and terrestrial plants; intermediate levels included aquatic macroinvertebrates, insects, and fish and bird guilds; and top predators included carnivorous fishes and aquatic predatory birds. These networks were used to identify dominant energy pathways and provide trophic context for interpreting Hg biomagnification across aquatic and riparian food webs.

## Results

### Trophic guild classification

The analyses included 132 consumers: 107 fishes and 25 birds. Fishes were classified as invertivores (n = 36), primary consumers (n = 27), carnivore–piscivores (n = 28), or omnivores (n = 16; Table S1). Carnivore–piscivores, including *Raphiodon vulpinus*, *Potamotrygon motoro*, *Pristobrycon calmoni*, and *Serrasalmus* spp., consumed predominantly fish prey. Invertivores and omnivores relied more strongly on aquatic macroinvertebrates, insects, detritus, and mixed resources.

Hierarchical clustering supported these assignments by separating carnivore–piscivores from primary consumers and detritivorous taxa, with intermediate consumers occupying transitional positions (Figure S1). This organization reflected differences in diet, isotopic composition, THg concentration, and body size.

Birds were classified as invertivores (n = 8), nectarivores (n = 8), frugi-granivores (n = 4), omnivores (n = 3), or aquatic predators (n = 2; Tables S1–S2). These guilds differed in their dominant resources, foraging ecology, and association with aquatic and riparian habitats.

### Mercury biomagnification

THg concentrations increased significantly with δ¹ N in fishes and birds, supporting trophic biomagnification (Figure 2). In fishes, δ¹ N explained 22.9% of THg variation (R² = 0.229, adjusted R² = 0.221; F = 30.29, p < 0.001). Carnivore–piscivores generally exhibited the highest δ¹ N and THg values, whereas primary consumers occupied lower positions along both gradients.

**Figure 2.**
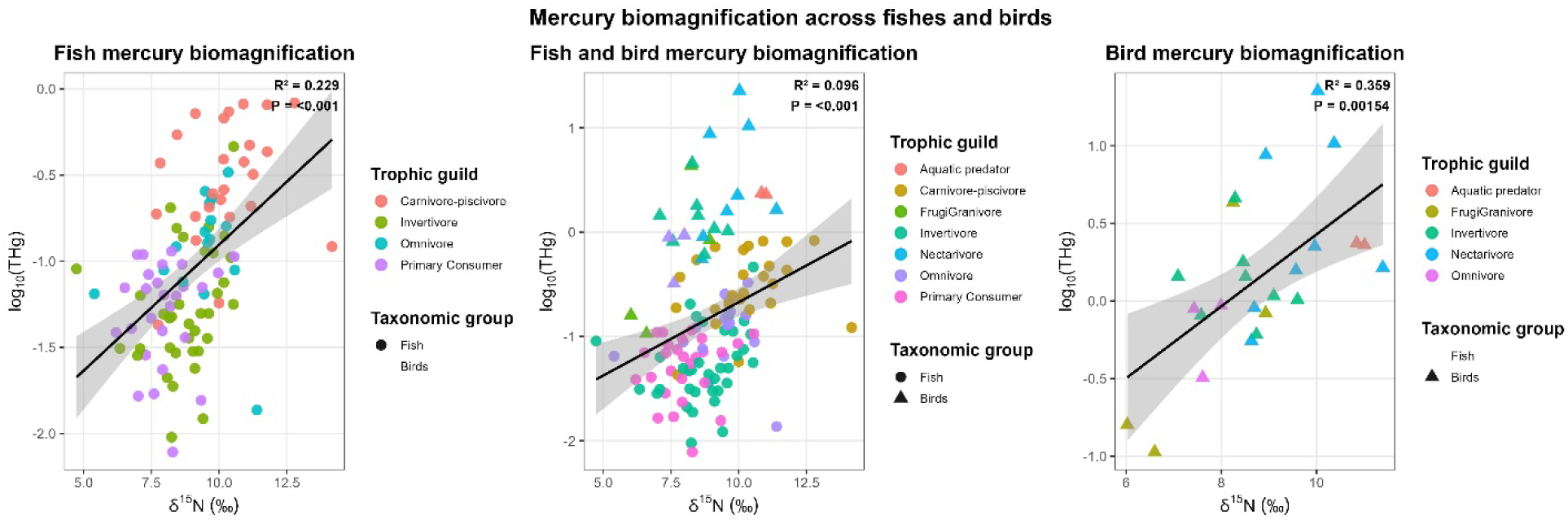
Relationships between δ¹ N and log□□-transformed total mercury concentrations for (A) fishes, (B) fishes and birds combined, and (C) birds. Colors indicate trophic guilds, symbols distinguish taxonomic groups, solid lines represent fitted linear models, and shaded areas indicate 95% confidence intervals.

Birds showed a stronger relationship, with trophic position explaining 35.9% of THg variation (R² = 0.359, adjusted R² = 0.332; F = 12.90, p = 0.0015). Aquatic predators exhibited higher THg concentrations than omnivorous and frugi-granivorous birds, consistent with exposure through aquatic prey.

Nectarivorous birds also showed unexpectedly high THg concentrations. Because this guild comprised hummingbirds rather than aquatic predators, this result should not be interpreted as evidence of a nectar-based Hg pathway. Instead, it may reflect arthropod supplementation, feather integration, or exposure across aquatic–riparian interfaces.

In the combined fish–bird dataset, THg remained positively associated with δ¹ N (R² = 0.096, adjusted R² = 0.089; F = 13.52, p < 0.001). Although the model explained less variation, the relationship indicated a consistent increase in Hg across trophic levels and taxonomic groups.

### Bayesian model performance

All MixSIAR models showed satisfactory MCMC convergence. Maximum Gelman–Rubin potential scale reduction factors ranged from 1.001 for aquatic predator birds to 1.034 for omnivorous birds and remained below the 1.05 threshold.

Geweke diagnostics indicated good stationarity for aquatic predator and frugi-granivorous birds and for primary consumer and invertivorous fishes. More outliers occurred in omnivorous and carnivore–piscivorous fishes and in omnivorous and nectarivorous birds, particularly in Chain 3. Nevertheless, the low PSRF values supported the overall reliability of posterior estimates, although results from the models with weaker Geweke performance should be interpreted more cautiously.

### Isotopic mixing spaces

Isotopic mixing spaces differentiated trophic guilds and their candidate resources (Figure 3). Primary consumer fishes occupied the most depleted positions (δ¹³C: approximately −37.9 to −22.5‰; δ¹ N: 6.2–10.5‰) and overlapped mainly with phytoplankton, detritus, and macrophytes. Periphyton was comparatively enriched and isotopically distinct.

**Figure 3.**
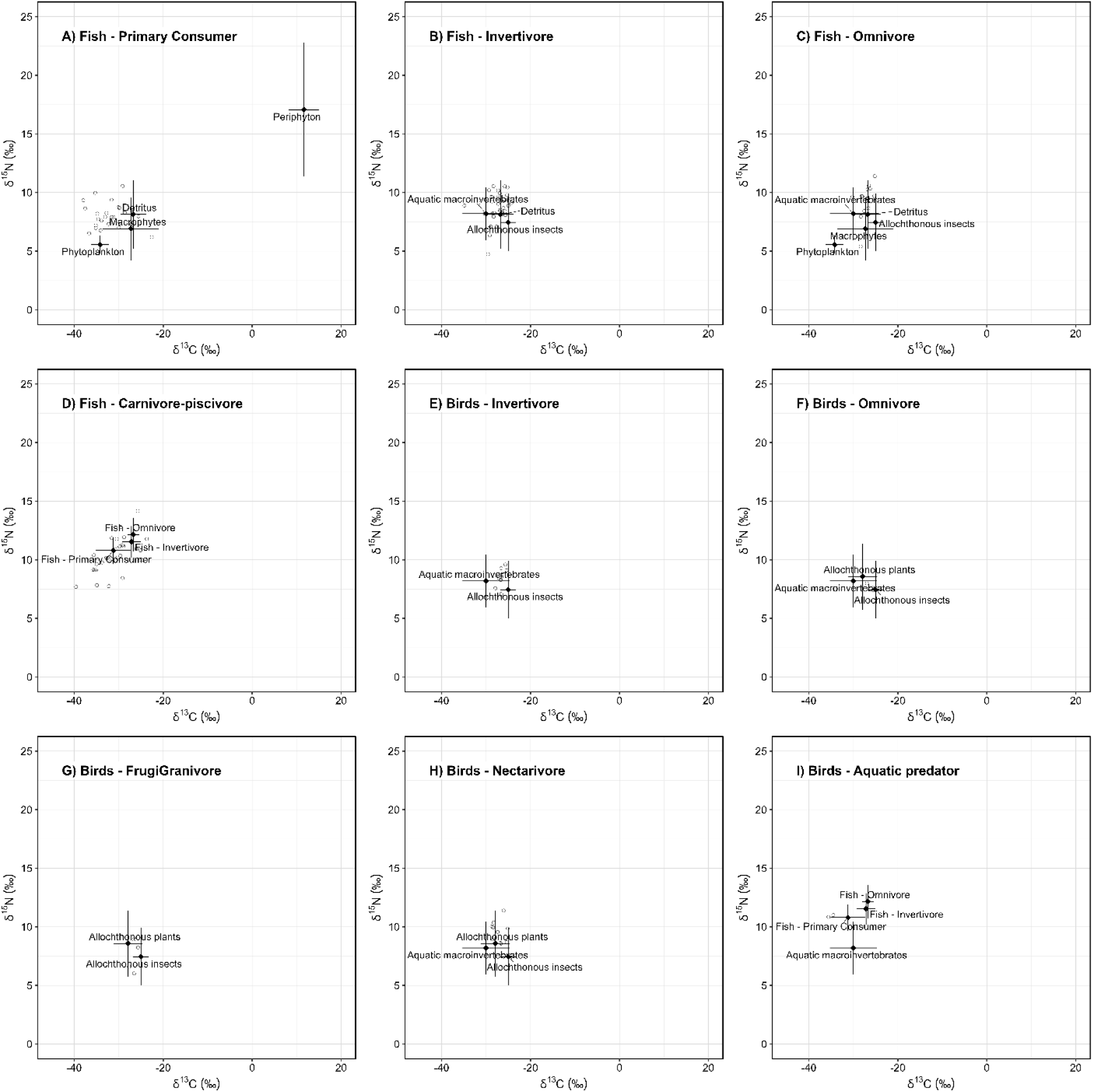
Isotopic mixing spaces used in Bayesian models for (A) fish primary consumers, (B) fish invertivores, (C) fish omnivores, (D) fish carnivore–piscivores, (E) bird invertivores, (F) bird omnivores, (G) bird frugi-granivores, (H) bird nectarivores, and (I) aquatic predator birds. Open circles represent consumers, while black symbols and error bars show source means ± SD after applying trophic enrichment factors.

Invertivorous fishes occupied intermediate positions (δ¹³C: approximately −32 to −24‰; δ¹ N: 8–11‰), overlapping with aquatic macroinvertebrates, allochthonous insects, and detritus. Carnivore–piscivores exhibited the broadest isotopic range (δ¹³C: −39.60 to −23.69‰; δ¹ N: 7.69–14.17‰), indicating heterogeneous basal carbon sources and their position near the upper end of the aquatic trophic gradient.

Bird guilds also exhibited distinct isotopic patterns. Invertivores clustered near allochthonous insects and aquatic macroinvertebrates, whereas frugi-granivores were closer to terrestrial plant resources. Omnivores showed the broadest avian carbon niche, consistent with the assimilation of diverse resources. Nectarivores exhibited enriched δ¹ N values despite their primarily plant-associated classification, supporting the possible importance of arthropod supplementation. Aquatic predators occupied the highest avian trophic positions and overlapped with fish-based pathways, although their small sample size warrants caution.

### Resource contributions

Bayesian models identified distinct trophic pathways among consumer guilds (Table 2). Primary consumer fishes were supported mainly by phytoplankton (62.5%), followed by detritus (22.5%), with smaller contributions from macrophytes and periphyton.

**Table 1.**
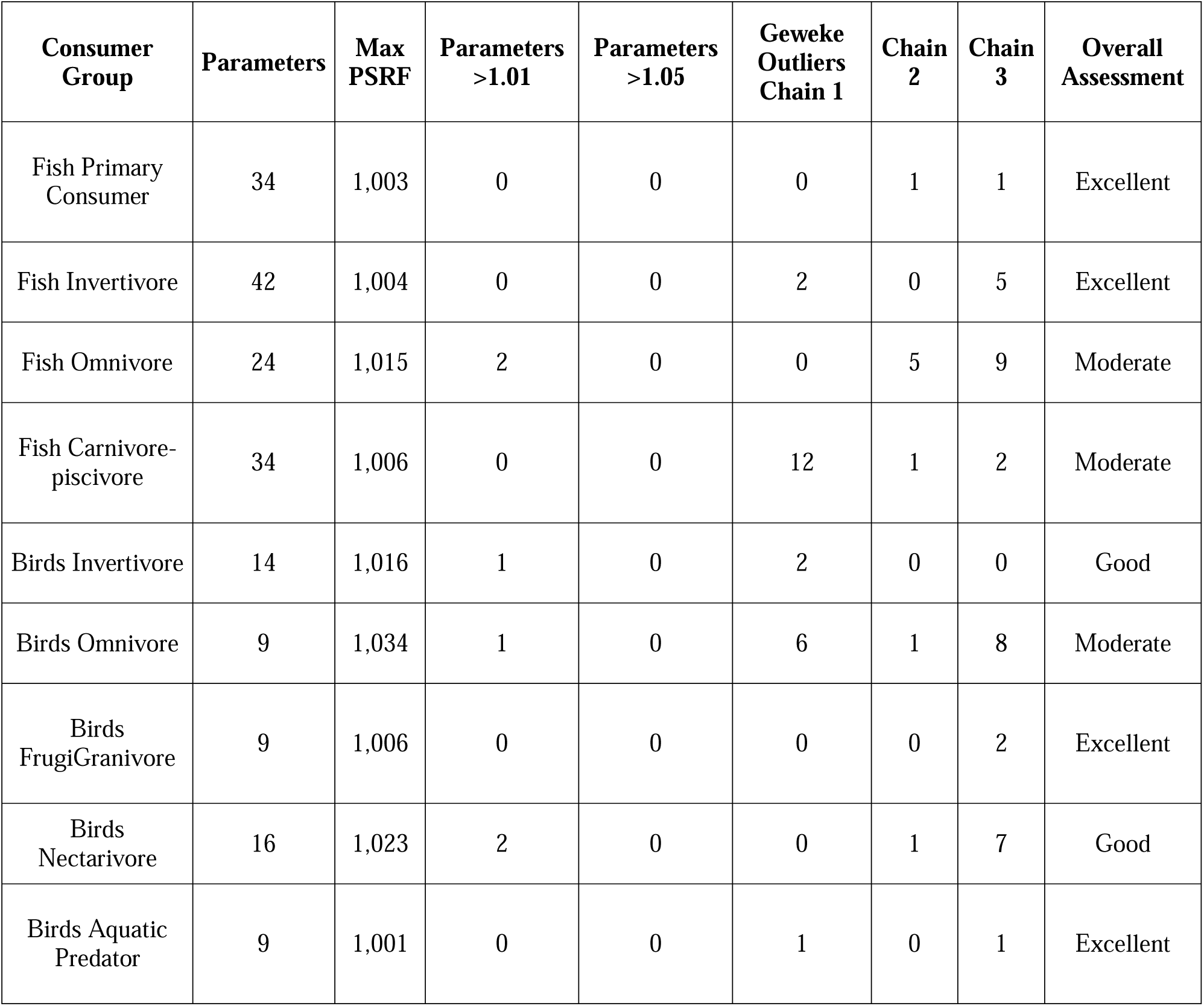
Convergence diagnostics for MixSIAR models, including maximum Gelman–Rubin potential scale reduction factors, parameters exceeding convergence thresholds, and Geweke outliers for each MCMC chain.

| Consumer Group | Parameters | Max PSRF | Parameters >1.01 | Parameters >1.05 | Geweke Outliers Chain 1 | Chain 2 | Chain 3 | Overall Assessment |
| --- | --- | --- | --- | --- | --- | --- | --- | --- |
| Fish Primary Consumer | 34 | 1,003 | 0 | 0 | 0 | 1 | 1 | Excellent |
| Fish Invertivore | 42 | 1,004 | 0 | 0 | 2 | 0 | 5 | Excellent |
| Fish Omnivore | 24 | 1,015 | 2 | 0 | 0 | 5 | 9 | Moderate |
| Fish Carnivore-piscivore | 34 | 1,006 | 0 | 0 | 12 | 1 | 2 | Moderate |
| Birds Invertivore | 14 | 1,016 | 1 | 0 | 2 | 0 | 0 | Good |
| Birds Omnivore | 9 | 1,034 | 1 | 0 | 6 | 1 | 8 | Moderate |
| Birds FrugiGranivore | 9 | 1,006 | 0 | 0 | 0 | 0 | 2 | Excellent |
| Birds Nectarivore | 16 | 1,023 | 2 | 0 | 0 | 1 | 7 | Good |
| Birds Aquatic Predator | 9 | 1,001 | 0 | 0 | 1 | 0 | 1 | Excellent |

**Table 2.**
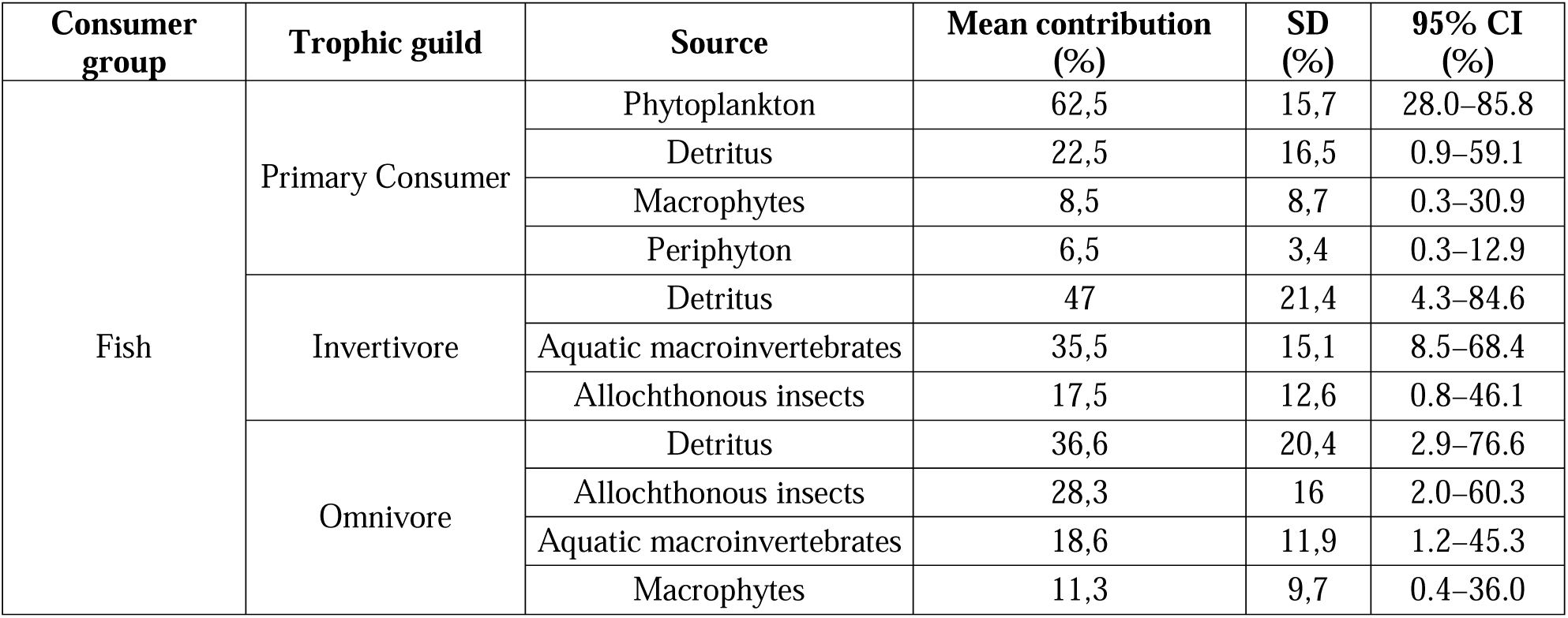
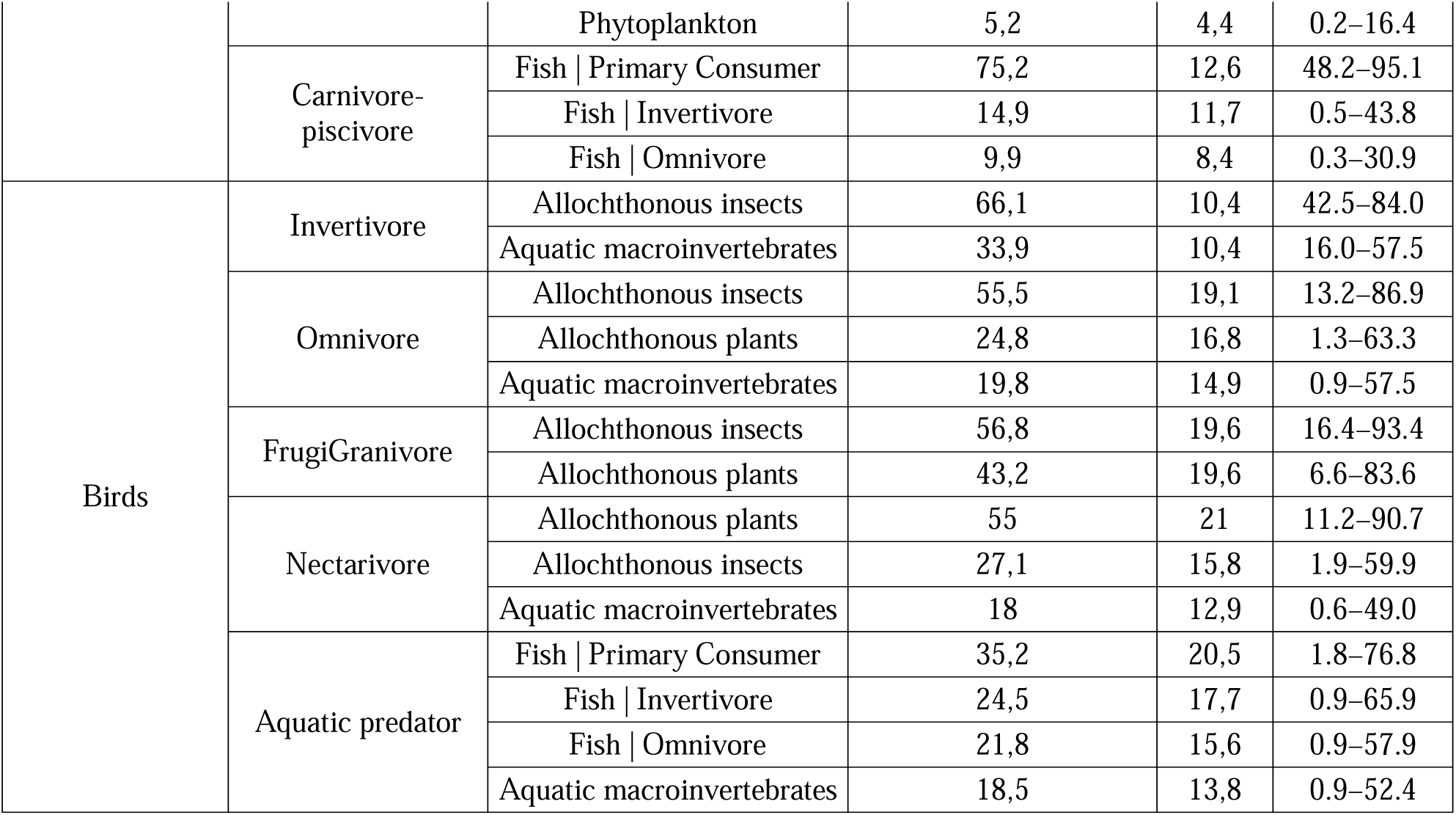
Posterior estimates of source contributions to fish and bird trophic guilds obtained from MixSIAR. Values represent posterior means, standard deviations, and 95% Bayesian credible intervals.

Invertivorous fishes relied primarily on detritus (47.0%) and aquatic macroinvertebrates (35.5%), with a smaller contribution from allochthonous insects (17.5%). Omnivorous fishes integrated detritus, insects, aquatic macroinvertebrates, macrophytes, and phytoplankton, although detritus remained dominant. Carnivore–piscivores relied strongly on primary consumer fishes (75.2%), with lower contributions from invertivorous and omnivorous fishes.

Among birds, invertivores depended primarily on allochthonous insects (66.1%). Omnivores also relied mainly on insects (55.5%), supplemented by terrestrial plants and aquatic macroinvertebrates. Frugi-granivores showed similar contributions from terrestrial plants and insects, whereas nectarivores were primarily supported by terrestrial plants (55.0%).

Aquatic predator birds integrated several aquatic pathways. Primary consumer fishes contributed 35.2%, followed by invertivorous fishes (24.5%), omnivorous fishes (21.8%), and aquatic macroinvertebrates (18.5%), indicating a relatively generalized feeding strategy.

### Trophic pathway reconstruction

The reconstructed network identified phytoplankton and detritus as the principal basal energy sources, with primary consumer fishes acting as the main link between basal production and upper-level consumers (Figure 4). The strongest pathway connected phytoplankton to primary consumer fishes and subsequently to carnivore–piscivores, which obtained 75.2% of their assimilated resources from primary consumer fishes.

**Figure 4.**
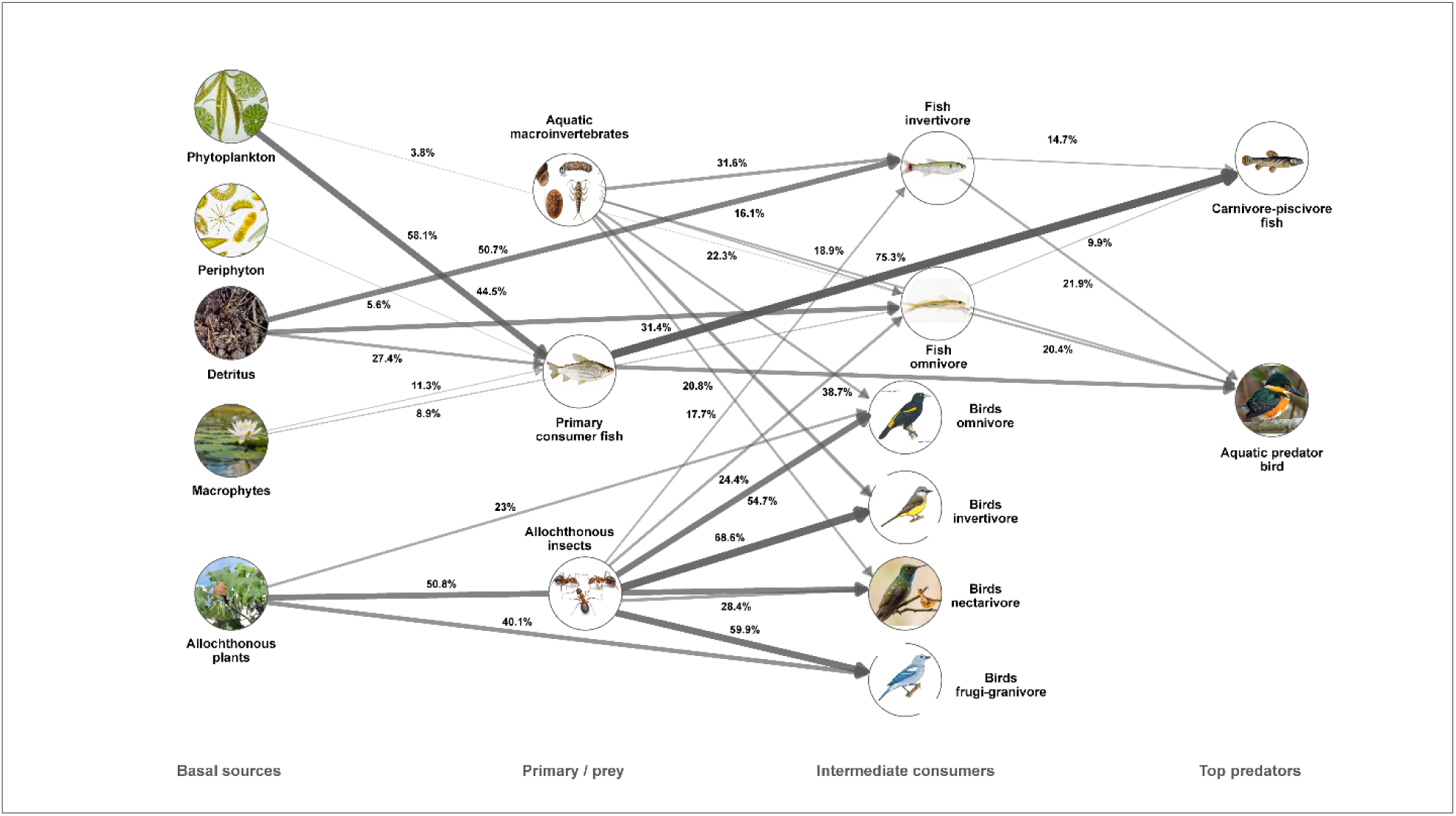
Bayesian trophic network reconstructed from MixSIAR posterior estimates. Nodes represent basal resources and consumer guilds, arrows indicate trophic pathways, and edge thickness is proportional to mean source contribution. Only contributions ≥10% are shown.

Primary consumer fishes also linked phytoplankton production to aquatic predator birds. Additional pathways through aquatic macroinvertebrates and invertivorous fishes further supported avian predators, demonstrating the contribution of aquatic invertebrate production to upper trophic levels. Terrestrial plants and allochthonous insects primarily supported non-predatory bird guilds.

Overall, phytoplankton- and detritus-based pathways supported intermediate fish consumers that subsequently transferred energy—and potentially Hg—to carnivore–piscivorous fishes and aquatic predator birds.

## Discussion

Our results show that mercury transfer in the upper Caquetá River basin is structured by trophic position and the organization of aquatic and aquatic-associated energy pathways. THg concentrations increased with δ¹ N in fishes, birds, and the combined fish–bird dataset, supporting trophic biomagnification. Bayesian mixing models further revealed that Hg exposure is embedded within a complex food web supported by phytoplankton, detritus, aquatic macroinvertebrates, terrestrial subsidies, and fish prey. Together, these findings indicate that Hg dynamics in this Amazonian river system are an emergent property of food-web structure rather than solely a species-level phenomenon.

### Trophic structure and mercury biomagnification in the upper Caquetá food web

The positive relationship between δ¹ N and log-transformed THg supports our first hypothesis: Hg biomagnifies along the trophic gradient of the upper Caquetá food web. This pattern was evident in fishes, birds, and the pooled dataset, indicating that organisms occupying higher trophic positions accumulate greater Hg burdens. This result is consistent with the behavior of methylmercury in aquatic food webs, where efficient assimilation, strong protein binding, and slow elimination promote progressive accumulation from prey to predators (Ullrich et al., 2001; Coelho et al., 2013; Lavoie et al., 2013).

In fishes, δ¹ N explained a substantial proportion of THg variation, with carnivore–piscivorous species occupying the upper isotopic and mercury gradients, particularly *Zungaro zungaro*, *Brachyplatystoma vaillantii*, *Pseudoplatystoma fasciatum*, *Raphiodon vulpinus*, and *Plagioscion squamosissimus*. This agrees with previous studies showing that Amazonian piscivores accumulate high Hg concentrations by integrating exposure across several prey species and trophic transfers (Barbosa et al., 2003; Pouilly et al., 2013; Olivero-Verbel et al., 2016; Mussy et al., 2023). Large, long-lived migratory predators such as *Z. zungaro*, *B. vaillantii*, and *P. fasciatum* may integrate contamination across multiple habitats by consuming fish prey, including migratory detritivores. Their Hg burdens therefore reflect both local exposure and broader ecological processes connecting migration, productivity, and contaminant transfer. Our results extend previous evidence of Hg contamination and human exposure risk in the Caquetá River (Olivero-Verbel et al., 2016) by demonstrating that accumulation is systematically structured by trophic position and food-web pathways.

The avian dataset also showed a positive and significant relationship between δ¹ N and THg, suggesting that Hg biomagnification extends beyond strictly aquatic consumers to riparian and aquatic-associated birds. This is important because tropical freshwater Hg studies have focused primarily on fish, while birds remain comparatively underrepresented despite their value as contamination sentinels (Evers et al., 2005; Clayden et al., 2015; Sayers et al., 2023). Aquatic predatory birds occupied high trophic positions and exhibited elevated THg relative to lower-trophic guilds, consistent with their consumption of fish and aquatic macroinvertebrates. Similar patterns have been reported in Neotropical mining-impacted landscapes, where piscivorous and insectivorous birds may accumulate substantial Hg burdens (Sierra-Marquez et al., 2018; Sayers et al., 2023; Pisconte et al., 2024).

The lower explanatory power of the pooled fish–bird model, relative to the separate models, indicates that the relationship between trophic position and Hg accumulation is not homogeneous across taxonomic groups. This likely reflects differences in dietary specialization, habitat use, mobility, lifespan, metabolism, and exposure pathways. Fishes and birds also differ in physiology, tissue type, exposure window, movement ecology, and Hg elimination (Matos et al., 2025). Fish muscle generally reflects relatively long-term aquatic exposure and local food-web integration, whereas feathers record Hg incorporated during feather growth and may integrate exposure across the period and location of molt (Evers et al., 2005; Sayers et al., 2023). Therefore, the pooled analysis demonstrates a broad trophic signal across the aquatic–terrestrial interface but should not be interpreted as evidence of a universal biomagnification slope shared by all consumers.

The moderate R² values further indicate that trophic position is an important but incomplete predictor of Hg concentrations. This agrees with global syntheses showing substantial variation in biomagnification slopes among ecosystems, even under comparable isotopic frameworks (Lavoie et al., 2013). Such variability may result from differences in methylmercury production, hydrological connectivity, sediment dynamics, primary productivity, food-chain length, growth, body size, age structure, and species-specific physiology (Ullrich et al., 2001; Lavoie et al., 2013; Karimi et al., 2016; Mussy et al., 2023).

Many of these sources of variation are likely amplified by the spatial and temporal heterogeneity of Amazonian river–floodplain systems. The study area includes a whitewater main channel, blackwater tributaries, and floodplain environments that differ in water chemistry, organic matter dynamics, and habitat structure (Sioli, 1984; Junk et al., 2011). During the high-water season, flooding increases connectivity among channels, flooded forests, lakes, and riparian habitats, creating environmental mosaics that influence Hg methylation and exposure pathways (Junk et al., 1989; Diringer et al., 2015). Riparian deforestation and land-use change in areas such as Curillo and San Antonio may further alter floodplain habitats and promote the mobilization of Hg and organic matter. Thus, although the biomagnification pattern supports H1, the unexplained variation suggests that Hg accumulation also depends on habitat-specific methylation, hydrology, and the carbon pathways supporting consumers.

### Basal-resource pathways and trophic routes of mercury transfer

Our second hypothesis proposed that Hg transfer would be influenced by basal-resource pathways rather than exclusively by trophic position. The results support this hypothesis because δ¹³C and δ¹ N provide complementary information on basal-resource use and trophic enrichment, respectively, while Bayesian mixing models estimate the proportional contribution of sources to consumer biomass (Peterson & Fry, 1987; Vander Zanden & Rasmussen, 2001; Post, 2002; Stock et al., 2018). These models identified dominant energy pathways connecting basal resources, fish guilds, and aquatic or riparian birds. However, the reconstructed pathways do not represent direct measurements of Hg flux. Instead, they provide the trophic context for interpreting biomagnification because Hg accumulation is mediated by the dietary routes through which methylmercury moves from basal resources and prey to upper consumers (Riva-Murray et al., 2013; Lavoie et al., 2013; Chen et al., 2014; Karimi et al., 2016).

The reconstructed pathways indicate that phytoplankton- and detritus-derived resources contribute substantially to the aquatic food web of the upper Caquetá basin. Primary consumer fishes showed a high probability of assimilating phytoplankton-derived carbon, with detritus representing a secondary pathway. Invertivorous fishes relied more strongly on detrital matter and aquatic macroinvertebrates, whereas omnivores integrated a broader range of resources. Hg transfer is therefore embedded within a food web combining pelagic, detrital, benthic-associated, and terrestrial subsidy pathways rather than following a single linear route. This structure is consistent with Amazonian rivers, where hydrological connectivity and seasonal flooding promote interactions among aquatic production, detritus, floodplain vegetation, and allochthonous inputs (Forsberg et al., 1993; Jepsen & Winemiller, 2002; Laffont et al., 2021).

The strongest reconstructed route was phytoplankton → primary consumer fishes → carnivore–piscivorous fishes (Figure 4; Table 2). This result indicates that upper-level fish predators are closely connected to prey assimilating autochthonous aquatic production, consistent with the importance of aquatic primary producers in Amazonian and tropical river food webs (Forsberg et al., 1993; Jepsen & Winemiller, 2002). Accordingly, Hg exposure in piscivorous fishes may be influenced not only by benthic or detrital processes but also by pathways supported by suspended aquatic production, as biomagnification depends on the routes through which methylmercury is transferred from lower trophic levels to predators (Lavoie et al., 2013; Chen et al., 2014; Karimi et al., 2016).

Detritus and aquatic macroinvertebrates also formed important pathways to invertivorous and omnivorous fishes, while macroinvertebrates contributed to aquatic predatory birds. Hg may therefore reach upper consumers through several aquatic and aquatic-associated channels, including phytoplankton-based, detrital, macroinvertebrate-mediated, and fish-prey pathways. Such multiplicity is expected in tropical rivers, where consumers integrate resources from several habitats rather than feeding within strictly compartmentalized pelagic or benthic chains (Forsberg et al., 1993; Jepsen & Winemiller, 2002; Laffont et al., 2021). It is also consistent with evidence that pelagic, benthic, detrital, and prey-mediated pathways differ in their contribution to Hg exposure (Riva-Murray et al., 2013; Chen et al., 2014; Karimi et al., 2016; Swinton et al., 2022).

These findings agree with evidence that basal-resource pathways influence Hg exposure. Methylation is frequently enhanced in organic-rich sediments, flooded soils, periphyton, and detrital habitats, from which lower-trophic consumers transfer methylmercury to fish and wildlife (Ullrich et al., 2001; Achá et al., 2011; Chen et al., 2014; Paranjape & Hall, 2017). Benthic and pelagic prey may therefore contribute differently to fish Hg burdens, while carbon sources can explain variation beyond trophic position (Riva-Murray et al., 2013; Karimi et al., 2016; Swinton et al., 2022). Nevertheless, the upper Caquetá food web cannot be reduced to a benthic–pelagic dichotomy: phytoplankton supported an important pathway to fish predators, while detritus and macroinvertebrates sustained intermediate consumers and aquatic-associated birds.

Therefore, H2 is supported as a food-web pathway hypothesis. The isotopic models identify energy flows and plausible routes through which Hg exposure may propagate, but they do not quantify how much Hg is associated with each source or pathway. Consumer concentrations also depend on source-specific methylmercury content, methylation rates, habitat exposure, and physiology. Future analyses should incorporate δ¹³C, δ¹ N, trophic guild, taxonomic group, locality, and their interactions as predictors of log-transformed THg. Direct methylmercury measurements in sediments, basal resources, periphyton, macroinvertebrates, and fish prey would further connect the reconstructed energy pathways with contaminant concentrations.

### Fish and aquatic birds as connected upper consumers: evidence for cross-boundary mercury transfer

Our third hypothesis predicted that upper-level consumers, particularly piscivorous fishes and aquatic birds, would integrate the highest Hg burdens and serve as sentinels of ecosystem contamination. The results broadly support this hypothesis, although the strength of inference differs between fishes and birds. Evidence was stronger for fishes because carnivore–piscivores were well represented and exhibited both enriched δ¹ N and elevated THg. They also formed a distinct trophic group and were strongly supported by primary consumer fishes in the mixing models. Piscivorous fishes are therefore important endpoints of aquatic biomagnification and remain central to ecological monitoring and human exposure assessments in the Caquetá basin.

For birds, the evidence was ecologically meaningful but limited by sample size. Aquatic predators occupied high trophic positions and exhibited elevated Hg concentrations, while mixing models indicated that they assimilated resources from primary consumer, invertivorous, and omnivorous fishes, as well as aquatic macroinvertebrates. This generalized resource use suggests exposure through several aquatic prey pathways rather than a single prey type. Such integration is consistent with the role of birds as sentinels because they connect aquatic food webs with riparian and terrestrial habitats and may reflect exposure across broader spatial and temporal scales than many fishes (Evers et al., 2005; Clayden et al., 2015; Sayers et al., 2023).

Nevertheless, the aquatic predator guild included only two individuals, so these findings constitute preliminary evidence rather than a definitive test of avian sentinel performance. The pattern supports including aquatic and riparian birds in Hg monitoring but cannot be generalized across species, habitats, or seasons. Neotropical avian Hg exposure may vary among trophic guilds, foraging strata, habitat associations, molt strategies, and levels of mining influence (Sayers et al., 2023; Pisconte et al., 2024). Larger samples of aquatic predators, riparian insectivores, and other aquatic-associated birds are needed to determine whether avian guilds reflect local aquatic pathways or broader landscape-level exposure.

Non-aquatic and partially aquatic-associated guilds provided an important ecological contrast. Invertivorous birds were supported primarily by allochthonous insects, whereas frugi-granivores and nectarivores were more closely linked to plant-derived resources. These guilds should not be considered direct indicators of aquatic biomagnification in the same way as aquatic predators, but they help distinguish consumers connected to aquatic prey from those primarily supported by terrestrial resources.

The unexpectedly elevated THg concentrations in nectarivorous birds, all of which were hummingbirds, require caution. They should not be interpreted as evidence of nectar-mediated Hg exposure because nectarivory describes the dominant energetic resource rather than the complete trophic niche. Hummingbirds supplement nectar with arthropods to meet protein and nitrogen requirements, potentially increasing their trophic position (Hardesty, 2009; Wilman et al., 2014; Billerman et al., 2026). Accordingly, their enriched δ¹ N values suggest some assimilation of animal resources. Elevated THg may therefore reflect arthropod-mediated, riparian, or landscape-level exposure rather than direct aquatic predation. Because the present study was not designed to distinguish these mechanisms, this result should be considered hypothesis-generating and evaluated through targeted sampling.

In Amazonian riparian landscapes, Hg exposure may arise through aquatic prey consumption and terrestrial pathways influenced by atmospheric deposition, canopy retention, and the movement of Hg across ecosystem boundaries via emerging aquatic insects (Evers et al., 2005; Rimmer et al., 2010; Wang et al., 2016; Gerson et al., 2017; Jackson et al., 2021; Twining et al., 2021). Although the present dataset cannot separate these mechanisms, it highlights the value of sampling birds from contrasting trophic guilds when investigating Hg movement across aquatic–terrestrial interfaces.

Fishes and birds thus provide complementary information. Fishes are directly connected to aquatic Hg dynamics and are essential for evaluating biomagnification and human consumption risk. Birds, particularly aquatic predators and riparian insectivores, can indicate whether Hg moves into mobile vertebrates integrating several habitats. This combined approach is especially relevant in the upper Caquetá, where Hg contamination associated with gold mining interacts with complex floodplain and riparian linkages. Consequently, H3 is strongly supported for piscivorous fishes and cautiously supported for aquatic predatory birds, which remain a promising but undersampled component of Hg biomonitoring.

### Ecological interpretation, limitations, and future directions

The upper Caquetá basin provides an important setting for studying Hg transfer because it combines mining influence, high rainfall, floodplain connectivity, and diverse aquatic and riparian food webs (Olivero-Verbel et al., 2016; Nogales et al., 2023). Artisanal and small-scale gold mining has occurred in the basin for decades, and Hg contamination has been documented in fish and human populations in the Colombian Amazon (Olivero-Verbel et al., 2016). Our findings indicate that exposure is not restricted to isolated taxa but is organized along trophic pathways connecting basal resources, intermediate consumers, fish predators, and aquatic-associated birds. Because sampling occurred during the rainy season, flood-pulse connectivity may have increased exchanges of organic matter and prey among channels, floodplain lakes, and flooded forests, potentially enhancing both food-web complexity and Hg transfer (Junk et al., 1989).

Several limitations should be considered. First, sampling covered a single hydrological period. Methylation, resource availability, fish movement, floodplain connectivity, and prey composition vary seasonally; therefore, the results represent a rainy-season snapshot rather than a complete annual assessment. Sampling during both dry and wet seasons would determine whether trophic pathways and biomagnification slopes remain stable across hydrological phases.

Second, locality data were pooled in the mixing models because of limited sample sizes within several guilds. Although this enabled the reconstruction of regional pathways, it reduced the ability to resolve spatial variation associated with mining pressure, sediment dynamics, and habitat structure. Hierarchical or mixed-effects models could explicitly evaluate spatial variability in trophic organization and Hg accumulation.

Third, bird sample sizes were limited, particularly for aquatic predators, restricting generalization across species, habitats, and seasons. Elevated hummingbird THg should likewise be treated as exploratory rather than evidence of a defined nectarivore pathway. Feather Hg represents exposure during feather formation and may not correspond precisely to the sampling location because of movement and molt timing. Future studies should combine larger avian samples, species-specific natural history and molt information, repeated sampling, and direct analysis of arthropod prey.

Fourth, we measured THg rather than methylmercury. THg is a useful proxy, particularly in upper-level consumers where methylmercury often constitutes a large proportion of total Hg, but this proportion varies among tissues, taxa, and trophic levels. Direct methylmercury measurements in basal resources, macroinvertebrates, fish muscle, and bird feathers would provide a stronger mechanistic link between methylation environments and consumer exposure.

Fifth, Bayesian mixing models depend on source selection, isotopic separation, trophic enrichment factors, and model structure. Although convergence diagnostics generally supported posterior estimates, models with weaker Geweke performance require greater caution. Source contributions represent assimilated diet rather than direct contaminant flux (Chiaradia et al., 2014); therefore, the reconstructed network describes likely energy pathways, not a quantitative map of Hg transfer. Future studies could incorporate source-specific Hg concentrations into concentration-dependent models (Phillips & Koch, 2002), apply compound-specific isotope analysis of amino acids (McMahon & McCarthy, 2016), or integrate stomach contents and DNA metabarcoding as Bayesian priors (Chiaradia et al., 2014).

Finally, fish guild classification incorporated diet, isotopic signatures, body size, and THg. Although this integrative approach identified ecologically coherent guilds, guild-level differences in Hg are not fully independent of the classification process. Consequently, the strongest evidence for biomagnification is the continuous positive relationship between δ¹ N and log-transformed THg rather than contrasts among guilds. Recognizing this distinction avoids circular interpretation and grounds the conclusions in the most robust statistical evidence.

## Conclusions

This study shows that mercury biomagnifies across the fish and bird components of the upper Caquetá River food web. The positive relationships between δ¹ N and THg support trophic position as a major predictor of mercury accumulation, particularly among carnivore–piscivorous fishes and aquatic predatory birds. Bayesian mixing models further indicate that mercury exposure is embedded within a complex food web supported by phytoplankton, detritus, aquatic macroinvertebrates, terrestrial subsidies, and fish prey. The unexpectedly elevated THg values observed in hummingbirds further suggest that mercury exposure in riparian bird assemblages may extend beyond strictly piscivorous or aquatic predatory guilds, although this pattern requires targeted evaluation before its mechanisms can be established.

The evidence supports H1 strongly, supports H2 indirectly through reconstructed energy pathways, and supports H3 most clearly for piscivorous fishes and more cautiously for aquatic predatory birds. Overall, the upper Caquetá River basin appears to function as a connected aquatic–riparian system in which mercury contamination can move from basal resources and intermediate consumers toward fish predators and birds. Future work should combine seasonal sampling, site-specific models, larger avian datasets, and direct methylmercury measurements to clarify how hydrology, basal carbon pathways, and mining influence interact to shape mercury transfer in Amazonian food webs.

## Acknowledgements

We would like to express our sincere gratitude to all the people from the communities of Peñas Blancas, Curillo, and San Antonio for their invaluable support and companionship during the fieldwork. We especially thank Don Anoraldo, Dufay, Darwin, Victor, and Saulo, as well as the children who showed an interest in learning about aquatic ecosystems and contributed to making our field experiences especially meaningful. We are particularly grateful to Dr. Sebastián Reynaldi for his valuable contributions during fieldwork and for the insightful discussions and scientific advice that helped refine the research questions and strengthen the development of the study hypotheses. We are also grateful to the Universidad de la Amazonia and its team for their valuable support throughout this study. In particular, we thank Dr. Lis Manrique, Dr. Johannes Stevens Ramírez Carvajal, and Saray Karina Gualteros Esguerra for their support with the mercury analyses of the samples. We also sincerely thank the administrative team, especially Magda Luna and Anthony Martínez Aranzalez, for their invaluable assistance with the administrative processes associated with this research. We are deeply grateful to our entire research team for their dedication and commitment. We especially thank Yessica Gómez Canticus, our outstanding field and laboratory assistant, and the Universidad Nacional de Colombia, Sede Amazonia, for their support. We also thank Erica Valentina Ruiz Ayala and Lenny Stefan Agudelo Pachón, and Professor Clemencia Torres for her valuable support with the macroinvertebrate material from the Universidad de la Amazonia. We also thank David Felipe Bulla Guaqueta and the Universidad Nacional de Colombia, Sede Medellín, for providing access to the laboratory of the Department of Forest Sciences. Finally, we gratefully acknowledge Professor Vladimir Eliodoro Costa from the Centro de Isótopos Estáveis “Prof. Dr. Carlos Ducatti”, Instituto de Biociências de Botucatu, Universidade Estadual Paulista (UNESP), Campus Botucatu, Brazil, for his support with the stable isotope analyses.

## Data availability

Data and R scripts supporting the findings of this study are publicly available in Figshare at 10.6084/m9.figshare.34018671. Analysis scripts are also available through the associated GitHub repository at github.com/willo9303/mixsiar-reproducible.

## CRediT authorship contribution statement

William Gonzalez-Daza: Conceptualization, Methodology, Investigation, Software, Data curation, Formal analysis, Writing – original draft, Writing – review & editing, Visualization.

Angélica María Torres-Bejarano: Conceptualization, Methodology, Investigation, Data curation, Writing – original draft, Writing – review & editing.

Juan Camilo Ríos-Orjuela: Conceptualization, Methodology, Investigation, Data curation, Visualization, Writing – original draft, Writing – review & editing.

Santiago R. Duque: Conceptualization, Supervision, Project administration, Writing – review & editing, Funding acquisition.

## Ethics and permits

Field sampling and specimen handling were conducted under the Universidad de la Amazonia’s Framework Permit for the Collection of Specimens of Wild Species of Biological Diversity for Non-Commercial Scientific Research, granted by the Colombian National Environmental Licensing Authority (ANLA) through Resolution No. 01140 of 30 September 2016. The study was conducted under the authorized research project “Análisis del impacto socio ambiental del mercurio y tecnologías sostenibles para su remoción en la cuenca alta del río Caquetá-Caquetá-Putumayo” (project code 600.6.668). Export of biological samples to the Centro de Isótopos Estables Prof. Dr. Carlos Ducatti, Universidade Estadual Paulista (UNESP), Brazil, for laboratory analyses was authorized by ANLA under non-CITES export permit No. 003849, issued on 1 October 2024.

## Declaration of generative AI and AI-assisted technologies in the manuscript preparation process

During the preparation of this manuscript, the authors used ChatGPT (GPT-5.6 Sol; OpenAI) exclusively to assist with grammatical corrections, language editing, and textual review. All AI-assisted revisions were subsequently reviewed, verified, and, where necessary, modified by the authors, who take full responsibility for the final content of the manuscript. Generative AI was not used for data collection, data processing or analysis, statistical analyses, interpretation of results, generation of scientific conclusions, or any other research activity.

## References

1. Achá, D., Hintelmann, H., & Yee, J. (2011). Importance of sulfate-reducing bacteria in mercury methylation and demethylation in periphyton from the Bolivian Amazon region. Chemosphere, 82(6), 911–916.

2. Alcala-Orozco, M., Caballero-Gallardo, K., & Olivero-Verbel, J. (2019). Mercury exposure assessment in indigenous communities from Tarapaca village, Cotuhe and Putumayo Rivers, Colombian Amazon. Environmental Science and Pollution Research, 26(36), 36458–36467. 10.1007/s11356-019-06620-x

3. Alcala-Orozco, M., Caballero-Gallardo, K., & Olivero-Verbel, J. (2020). Biomonitoring of mercury, cadmium and selenium in fish and the population of Puerto Nariño, at the southern corner of the Colombian Amazon. Archives of Environmental Contamination and Toxicology, 79(3), 354–370. 10.1007/s00244-020-00761-8

4. Araruna, L. T., de Oliveira, A. T., de Oliveira Novaes, E., de Pinho, J. V., de Almeida Rodrigues, P., & Conte-Junior, C. A. (2025). Mercury contamination endangers Indigenous Amazonian communities: A systematic review. Environmental Science and Pollution Research, 32(23), 13607–13625.

5. Barbosa, A. C., Souza, J. D., Dórea, J. G., Jardim, W. F., & Fadini, P. S. (2003). Mercury biomagnification in a tropical black water, Rio Negro, Brazil. Archives of Environmental Contamination and Toxicology, 45(2), 235–246.

6. Bazhenova, D. E., Kapustina, V. Y., & Ivanova, E. S. (2024). Mercury content in the organs fish from different reservoirs of the Ustyuzhensky District of the Vologda Region. Samara Journal of Science, 13(4), 8–13. 10.55355/snv2024134101

7. Billerman, S. M., Keeney, B. K., Kirwan, G. M., Medrano, F., Sly, N. D., & Smith, M. G. (Eds.). (2026). Birds of the World. Cornell Laboratory of Ornithology. 10.2173/bow

8. Cardona, G. I., Escobar, M. C., Acosta-González, A., Marín, P., & Marqués, S. (2022). Highly mercury-resistant strains from different Colombian Amazon ecosystems affected by artisanal gold mining activities. Applied Microbiology and Biotechnology, 106(7), 2775–2793.

9. Casso-Hartmann, L., Vanegas, D. C., & McLamore, E. S. (2022). Water pollution and environmental policy in artisanal gold mining frontiers: The case of La Toma, Colombia. Science of the Total Environment, 852, 158417.

10. Chen, C. Y., Borsuk, M. E., Bugge, D. M., Hollweg, T., Balcom, P. H., Ward, D. M., & Sunderland, E. M. (2014). Benthic and pelagic pathways of methylmercury bioaccumulation in estuarine food webs. PLoS ONE, 9(2), e89305.

11. Chiaradia, A., Forero, M. G., McInnes, J. C., & Ramírez, F. (2014). Searching for the true diet of marine predators: Incorporating Bayesian priors into stable isotope mixing models. PLoS ONE, 9(3), e92665.

12. Clayden, M. G., Arsenault, L. M., Kidd, K. A., O’Driscoll, N. J., & Mallory, M. L. (2015). Mercury bioaccumulation and biomagnification in an Arctic marine food web. Science of the Total Environment, 509–510, 206–215.

13. Coelho, J. P., Mieiro, C. L., Pereira, E., Duarte, A. C., & Pardal, M. A. (2013). Mercury biomagnification in a contaminated estuary food web: Effects of age and trophic position using stable isotope analyses. Marine Pollution Bulletin, 69(1–2), 110–115.

14. Crespo-Lopez, M. E., Augusto-Oliveira, M., Lopes-Araújo, A., Santos-Sacramento, L., Takeda, P. Y., de Matos Macchi, B., & Arrifano, G. P. (2021). Mercury: What can we learn from the Amazon? Environment International, 146, 106223.

15. Diringer, S. E., Feingold, B. J., Ortiz, E. J., Gallis, J. A., Araújo-Flores, J. M., Berky, A., & Hsu-Kim, H. (2015). River transport of mercury from artisanal and small-scale gold mining and risks for dietary mercury exposure in Madre de Dios, Peru. Environmental Science: Processes & Impacts, 17(2), 478–487.

16. Evers, D. C., Burgess, N. M., Champoux, L., Hoskins, B., Major, A., Goodale, W. M., Taylor, R. J., Poppenga, R., & Daigle, T. (2005). Patterns and interpretation of mercury exposure in freshwater avian communities in northeastern North America. Ecotoxicology, 14(1–2), 193–221. 10.1007/s10646-004-6269-7

17. Forsberg, B. R., Araujo-Lima, C. A. R. M., Martinelli, L. A., Victoria, R. L., & Bonassi, J. A. (1993). Autotrophic carbon sources for fish of the central Amazon. Ecology, 74(3), 643–652.

18. Gasca Álvarez, A. D. P. (2000). Environmental exposure to mercury in gold mining: Health impact assessment in Guainía, Colombia. Revista de Salud Pública, 2(3), 233–250.

19. Gelman, A., Carlin, J. B., Stern, H. S., Dunson, D. B., Vehtari, A., & Rubin, D. B. (2014). Bayesian data analysis (3rd ed.). Chapman and Hall/CRC. 10.1201/b16018

20. Gerson, J. R., Driscoll, C. T., Demers, J. D., Sauer, A. K., Blackwell, B. D., Montesdeoca, M. R., Shanley, J. B., & Ross, D. S. (2017). Deposition of mercury in forests across a montane elevation gradient: Elevational and seasonal patterns in methylmercury inputs and production. Journal of Geophysical Research: Biogeosciences, 122(8), 1922–1939. 10.1002/2016JG003721

21. Geweke, J. (1991). Evaluating the accuracy of sampling-based approaches to the calculation of posterior moments (Staff Report No. 148). Federal Reserve Bank of Minneapolis, Research Department. 10.21034/sr.148

22. Gilmour, C. C., Podar, M., Bullock, A. L., Graham, A. M., Brown, S. D., Somenahally, A. C., & Elias, D. A. (2013). Mercury methylation by novel microorganisms from new environments. Environmental Science & Technology, 47(20), 11810–11820.

23. Guerrero, A. I., & Rogers, T. L. (2020). Evaluating the performance of the Bayesian mixing tool MixSIAR with fatty acid data for quantitative estimation of diet. Scientific Reports, 10(1), 20780. 10.1038/s41598-020-77396-1

24. Hardesty, J. L. (2009). Using nitrogen-15 to examine protein sources in hummingbird diets. Ornitología Colombiana, 8, 19–28.

25. Jackson, A. K., Eagles-Smith, C. A., & Robinson, W. D. (2021). Differential reliance on aquatic prey subsidies influences mercury exposure in riparian arachnids and songbirds. Ecology and Evolution, 11(11), 7003–7017. 10.1002/ece3.7549

26. Jepsen, D. B., & Winemiller, K. O. (2002). Structure of tropical river food webs revealed by stable isotope ratios. Oikos, 96(1), 46–55.

27. Junk, W. J., Bayley, P. B., & Sparks, R. E. (1989). The flood pulse concept in river-floodplain systems. In D. P. Dodge (Ed.), Proceedings of the International Large River Symposium (pp. 110–127). Canadian Special Publication of Fisheries and Aquatic Sciences, 106.

28. Junk, W. J., Piedade, M. T. F., Schöngart, J., Cohn-Haft, M., Adeney, J. M., & Wittmann, F. (2011). A classification of major naturally-occurring Amazonian lowland wetlands. Wetlands, 31(4), 623–640.

29. Karimi, R., Chen, C. Y., Folt, C. L., & Pickhardt, P. C. (2016). Comparing benthic and pelagic prey as mercury sources to fish. Science of the Total Environment, 565, 211–221.

30. Lacerda, L. D. (1997). Global mercury emissions from gold and silver mining. Water, Air, and Soil Pollution, 97(3), 209–221.

31. Laffont, L., Menges, J., Goix, S., Gentès, S., Maury-Brachet, R., Sonke, J. E., Legeay, A., Gonzalez, P., Rinaldo, R., & Maurice, L. (2021). Hg concentrations and stable isotope variations in tropical fish species of a gold-mining-impacted watershed in French Guiana. Environmental Science and Pollution Research, 28, 60609–60621.

32. Lavoie, R. A., Jardine, T. D., Chumchal, M. M., Kidd, K. A., & Campbell, L. M. (2013). Biomagnification of mercury in aquatic food webs: A worldwide meta-analysis. Environmental Science & Technology, 47(23), 13385–13394.

33. Lopes, L. E., Fernandes, A. M., Medeiros, M. C. I., & Marini, M. Â. (2016). A classification scheme for avian diet types. Journal of Field Ornithology, 87(3), 309–322. 10.1111/jofo.12158

34. Marrugo-Negrete, J., Benitez, L. N., & Olivero-Verbel, J. (2008). Distribution of mercury in several environmental compartments in an aquatic ecosystem impacted by gold mining in northern Colombia. Archives of Environmental Contamination and Toxicology, 55(2), 305–316.

35. Martoredjo, I., Calvão Santos, L. B., Vilhena, J. C. E., Rodrigues, A. B. L., de Almeida, A., Sousa Passos, C. J., & Florentino, A. C. (2024). Trends in mercury contamination distribution among human and animal populations in the Amazon region. Toxics, 12(3), 204.

36. Matos, L. S. D., Kasper, D., Silva, J. O. S., & Carvalho, L. N. (2025). Mercury in piscivorous and detritivorous fish from the Teles Pires river basin, Southern Amazon. Revista Brasileira de Ciências Ambientais, 60, e2144. 10.5327/Z2176-94782144

37. McDaniel, E. A., Peterson, B. D., Stevens, S. L. R., Tran, P. Q., Anantharaman, K., & McMahon, K. D. (2020). Expanded phylogenetic diversity and metabolic flexibility of mercury-methylating microorganisms. mSystems, 5(4), e00299–20. 10.1128/mSystems.00299-20

38. McMahon, K. W., & McCarthy, M. D. (2016). Embracing variability in amino acid δ¹ N fractionation: Mechanisms, implications, and applications for trophic ecology. Ecosphere, 7(12), e01511.

39. Meneses, H. do N. de M., Oliveira-da-Costa, M., Basta, P. C., Morais, C. G., Pereira, R. J. B., de Souza, S. M. S., & Hacon, S. de S. (2022). Mercury contamination: A growing threat to riverine and urban communities in the Brazilian Amazon. International Journal of Environmental Research and Public Health, 19(5), 2816. 10.3390/ijerph19052816

40. Mussy, M. H., De Almeida, R., De Carvalho, D. P., Lauthartte, L. C., De Holanda, I. B. B., Almeida, M. G. D., & Bastos, W. R. (2023). Evaluating mercury biomagnification using stable isotopes in Amazonian fish. Environmental Science and Pollution Research, 30(12), 33543–33554.

41. Nogales, J., Rogéliz-Prada, C., Cañon, M. A., & Vargas-Luna, A. (2023). An integrated methodological framework for the durable conservation of freshwater ecosystems: A case study in Colombia’s Caquetá River basin. Frontiers in Environmental Science, 11, 1264392.

42. Olivero-Verbel, J., Carranza-Lopez, L., Caballero-Gallardo, K., Ripoll-Arboleda, A., & Muñoz-Sosa, D. (2016). Human exposure and risk assessment associated with mercury pollution in the Caqueta River, Colombian Amazon. Environmental Science and Pollution Research, 23(20), 20761–20771.

43. Paiva, T. C., Pestana, I. A., de Oliveira, B. C. V., de Almeida, M. G., Malm, O., de Rezende, C. E., & Kasper, D. (2024). Mercury concentrations and differences in isotopic niches of fish from upstream and downstream of an Amazon reservoir dam. Ecotoxicology, 33(7), 762–771. 10.1007/s10646-024-02776-6

44. Paranjape, A. R., & Hall, B. D. (2017). Recent advances in the study of mercury methylation in aquatic systems. Facets, 2(1), 85–119.

45. Passos, C. J., & Mergler, D. (2008). Human mercury exposure and adverse health effects in the Amazon: A review. Cadernos de Saúde Pública, 24, s503–s520.

46. Peterson, B. J., & Fry, B. (1987). Stable isotopes in ecosystem studies. Annual Review of Ecology and Systematics, 18, 293–320.

47. Phillips, D. L., & Koch, P. L. (2002). Incorporating concentration dependence in stable isotope mixing models. Oecologia, 130(2), 114–125.

48. Pisconte, J. N., Vega, C. M., Sayers, C. J., Sevillano-Ríos, C. S., Pillaca, M., Quispe, E., Tejeda, V., Ascorra, C., Silman, M., & Fernandez, L. E. (2024). Elevated mercury exposure in bird communities inhabiting artisanal and small-scale gold mining landscapes of the southeastern Peruvian Amazon. Ecotoxicology, 33, 472–483.

49. Post, D. M. (2002). Using stable isotopes to estimate trophic position. Ecology, 83(3), 703–718.

50. Pouilly, M., Rejas, D., Pérez, T., Duprey, J. L., Molina, C. I., Hubas, C., & Guimarães, J. R. D. (2013). Trophic structure and mercury biomagnification in tropical fish assemblages, Iténez River, Bolivia. PLOS ONE, 8(5), e65054. 10.1371/journal.pone.0065054

51. Rimmer, C. C., Miller, E. K., McFarland, K. P., Taylor, R. J., & Faccio, S. D. (2010). Mercury bioaccumulation and trophic transfer in the terrestrial food web of a montane forest. Ecotoxicology, 19(4), 697–709. 10.1007/s10646-009-0443-x

52. Riva-Murray, K., Bradley, P. M., Chasar, L. C., Button, D. T., Brigham, M. E., Scudder Eikenberry, B. C., … & Lutz, M. A. (2013). Influence of dietary carbon on mercury bioaccumulation in streams of the Adirondack Mountains of New York and the Coastal Plain of South Carolina, USA. Ecotoxicology, 22(1), 60–71.

53. Rocha, A. R. M., Beneditto, A. P. M. D., Pestana, I. A., & Souza, C. M. M. (2015). Isotopic profile and mercury concentration in fish of the lower portion of the rio Paraíba do Sul watershed, southeastern Brazil. Neotropical Ichthyology, 13(3), 723–732. 10.1590/1982-0224-20150047

54. Rumiantseva, O., Ivanova, E., & Komov, V. (2022). High variability of mercury content in the hair of Russia Northwest population: The role of the environment and social factors. International Archives of Occupational and Environmental Health, 95(5), 1027–1042. 10.1007/s00420-021-01812-w

55. Sayers, C. J., Evers, D. C., Ruiz-Gutierrez, V., Adams, E., Vega, C. M., Pisconte, J. N., Tejeda, V., Regan, K., Lane, O., Ash, A., Cal, R., Reneau, S., Martínez, W., Welch, G., Hartwell, K., Teul, M., Tzul, D., Arendt, W., Tórrez, M., Watsa, M., Erkenswick, G., Moore, C., Gerson, J., Sánchez, V., Purizaca, R., Yurek, H., Burton, M., Shrum, P., Tabares-Segovia, S., Vargas, K., Fogarty, F., Charette, M., Martínez, A., Bernhardt, E., Taylor, R., Tear, T., & Fernandez, L. E. (2023). Mercury in Neotropical birds: A synthesis and prospectus on 13 years of exposure data. Ecotoxicology, 32(8), 1096–1123. 10.1007/s10646-023-02706-y

56. Selin, N. E. (2009). Global biogeochemical cycling of mercury: A review. Annual Review of Environment and Resources, 34, 43–63.

57. Sierra-Marquez, L., Peñuela-Gomez, S., Franco-Espinosa, L., Gomez-Ruiz, D., Diaz-Nieto, J., Sierra-Marquez, J., & Olivero-Verbel, J. (2018). Mercury levels in birds and small rodents from Las Orquideas National Natural Park, Colombia. Environmental Science and Pollution Research, 25(35), 35055–35063.

58. Sioli, H. (1984). The Amazon and its main affluents: Hydrography, morphology of the river courses, and river types. In H. Sioli (Ed.), The Amazon: Limnology and landscape ecology of a mighty tropical river and its basin (pp. 127–165). Springer.

59. Stock, B. C., Jackson, A. L., Ward, E. J., Parnell, A. C., Phillips, D. L., & Semmens, B. X. (2018). Analyzing mixing systems using a new generation of Bayesian tracer mixing models. PeerJ, 6, e5096. 10.7717/peerj.5096

60. Swinton, M. W., Eichler, L. W., & Nierzwicki-Bauer, S. A. (2022). Stable isotopes explain methylmercury concentrations in stream food webs. Ecotoxicology, 31, 1–14.

61. Terborgh, J., Robinson, S. K., Parker, T. A., III, Munn, C. A., & Pierpont, N. (1990). Structure and organization of an Amazonian forest bird community. Ecological Monographs, 60(2), 213–238. 10.2307/1943045

62. Twining, C. W., Razavi, N. R., Brenna, J. T., Dzielski, S. A., Gonzalez, S. T., Lawrence, P., Cleckner, L. B., & Flecker, A. S. (2021). Emergent freshwater insects serve as subsidies of methylmercury and beneficial fatty acids for riparian predators across an agricultural gradient. Environmental Science & Technology, 55(9), 5868–5877. 10.1021/acs.est.0c07683

63. Ullrich, S. M., Tanton, T. W., & Abdrashitova, S. A. (2001). Mercury in the aquatic environment: Factors affecting methylation. Critical Reviews in Environmental Science and Technology, 31(3), 241–293.

64. Vander Zanden, M. J., & Rasmussen, J. B. (2001). Variation in δ¹ N and δ¹³C trophic fractionation. Limnology and Oceanography, 46(8), 2061–2066.

65. Vargas Licona, S. P., & Marrugo Negrete, J. L. (2019). Mercurio, metilmercurio y otros metales pesados en peces de Colombia: Riesgo por ingesta. Acta Biológica Colombiana, 24(2), 232–242.

66. Veiga, M. M., & Hinton, J. J. (2002). Abandoned artisanal gold mines and mercury contamination. Journal of Cleaner Production, 10(1), 1–17.

67. Wang, X., Bao, Z., Lin, C.-J., Yuan, W., & Feng, X. (2016). Assessment of global mercury deposition through litterfall. Environmental Science & Technology, 50(16), 8548–8557. 10.1021/acs.est.5b06351

68. Wilman, H., Belmaker, J., Simpson, J., de la Rosa, C., Rivadeneira, M. M., & Jetz, W. (2014). EltonTraits 1.0: Species-level foraging attributes of the world’s birds and mammals. Ecology, 95(7), 2027. 10.1890/13-1917.1

