## Supplementary material for "Multiple energy pathways structure mercury biomagnification in an Amazonian River food web"

**Figure S1**. Hierarchical clustering of fish species based on dietary composition and trophic resource use. Species were grouped using a mixed trophic similarity matrix and average-linkage hierarchical clustering (UPGMA). Species connected at lower linkage heights exhibit more similar feeding strategies, whereas species joined at higher heights display greater trophic differentiation. The identified clusters were subsequently used to classify species into trophic guilds for Bayesian mixing models (MixSIAR) and mercury biomagnification analyses.

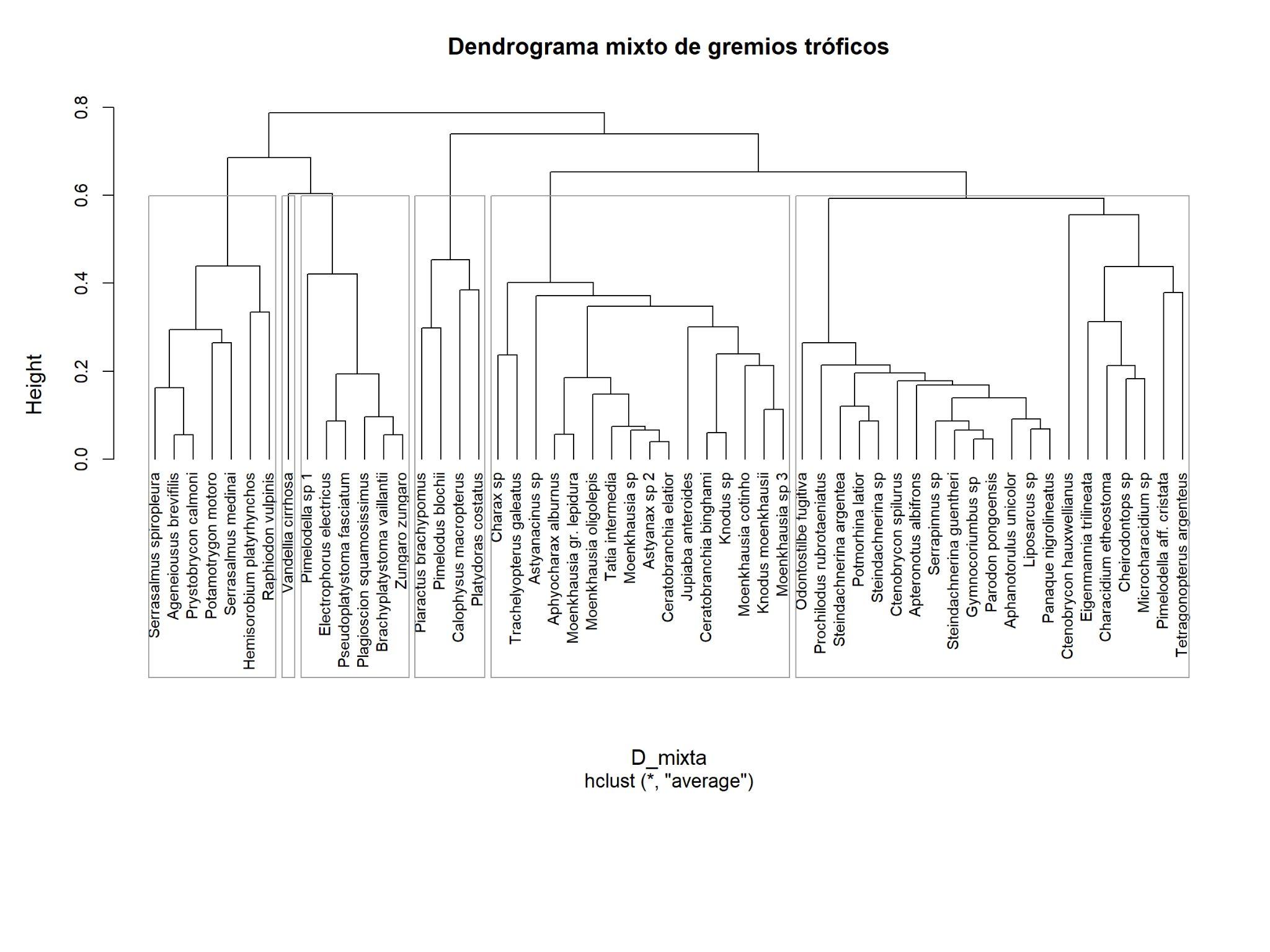

**Table S1.** Species-level summary of stable isotope signatures (δ¹³C and δ¹⁵N) and total mercury concentrations (THg) for fish and bird species sampled in the Colombian Amazon. Species are grouped according to their trophic guilds. Values are presented as mean ± standard deviation when more than one individual was available; for species represented by a single individual, the observed value is reported. Sample size (n) indicates the number of individuals analyzed per species. Total mercury (THg) concentrations are reported in mg kg⁻¹ dry weight. Stable isotope ratios are expressed in ‰ relative to international standards (Vienna Pee Dee Belemnite for carbon and atmospheric N₂ for nitrogen).

| **Group** | **Trophic guild** | **Species** | **n** | **δ13C mean ± SD** | **δ15N mean ± SD** | **THg mean ± SD** |
| --- | --- | --- | --- | --- | --- | --- |
| Birds | Aquatic predator | *Chloroceryle aenea* | 2 | -35.06 ± 0.76 | 10.93 ± 0.12 | 2.332 ± 0.049 |
|  | FrugiGranivore | *Leptotila rufaxilla* | 2 | -27.13 ± 0.81 | 6.31 ± 0.41 | 0.133 ± 0.038 |
|  | FrugiGranivore | *Thraupis episcopus* | 2 | -26.15 ± 0.69 | 8.59 ± 0.49 | 2.568 ± 2.449 |
|  | Invertivore | *Dendrexetastes rufigula* | 1 | -26.57 | 7.09 | 1.441 |
|  | Invertivore | *Dendrocolaptes picumnus* | 1 | -26.45 | 8.74 | 0.607 |
|  | Invertivore | *Dendrocyncla fuliginosa* | 1 | -26.57 | 8.51 | 1.433 |
|  | Invertivore | *Elaenia sp.* | 1 | -26.71 | 8.29 | 4.587 |
|  | Invertivore | *Pipra filicauda* | 1 | -27.91 | 7.58 | 0.811 |
|  | Invertivore | *Poecilotriccus latirostris* | 1 | -26.62 | 9.28 |  |
|  | Invertivore | *Tolmomyias sp.* | 1 | -27.14 | 8.46 | 1.783 |
|  | Invertivore | *Tyrannus melancholicus* | 1 | -25.66 | 9.6 | 1.019 |
|  | Invertivore | *Xiphorhynchus guttatus* | 1 | -25.4 | 9.1 | 1.078 |
|  | Nectarivore | *Amazilia versicolor* | 3 | -26.45 ± 1.18 | 9.35 ± 0.64 | 1.065 ± 0.722 |
|  | Nectarivore | *Florisuga mellivora* | 1 | -26.05 | 11.39 | 1.643 |
|  | Nectarivore | *Glaucis hirsutus* | 2 | -28.36 ± 0.08 | 10.16 ± 0.29 | 6.311 ± 5.74 |
|  | Nectarivore | *Phaethornis atrimentalis* | 1 | -28.73 | 10.03 | 22.6 |
|  | Nectarivore | *Phaethornis bourcieri* | 1 | -26.65 | 8.57 |  |
|  | Nectarivore | *Phaethornis hispidus* | 2 | -26.9 ± 2.15 | 8.81 ± 0.17 | 4.819 ± 5.534 |
|  | Omnivore | *Cacicus cela* | 2 | -25.58 ± 0.04 | 7.52 ± 0.13 | 0.607 ± 0.404 |
|  | Invertivore | *Monasa nigrifrons* | 1 | -26.95 | 7.99 | 0.934 |
| Fish | Carnivore-piscivore | *Ageneiousus brevifilis* | 1 | -30.32 | 10.18 | 0.26 |
|  | Carnivore-piscivore | *Brachyplatystoma vaillantii* | 1 | -31.6 | 11.86 | 2.278844 |
|  | Carnivore-piscivore | *Calophysus macropterus* | 2 | -25.57 ± 0.66 | 10.87 ± 0.08 | 0.377 |
|  | Carnivore-piscivore | *Electrophorus electricus* | 1 | -34.9 | 7.83 | 0.372 |
|  | Carnivore-piscivore | *Hemisorobium platyrhynchos* | 2 | -30.74 ± 1.3 | 10.6 ± 0.76 | 0.35 ± 0.173 |
|  | Carnivore-piscivore | *Hoplias malabaricus* | 4 | -32.07 ± 5.06 | 9.01 ± 0.93 | 0.104 ± 0.065 |
|  | Carnivore-piscivore | *Plagioscion squamosissimus* | 1 | -29.6 | 12.79 | 0.831 |
|  | Carnivore-piscivore | *Potamotrygon constellata* | 1 | -29.13 | 8.45 | 0.542 |
|  | Carnivore-piscivore | *Potamotrygon motoro* | 1 | -29.18 | 11.26 | 0.32 |
|  | Carnivore-piscivore | *Pristobrycon calmoni* | 1 | -32.52 | 10.17 | 0.391 |
|  | Carnivore-piscivore | *Pseudoplatystoma fasciatum* | 1 | -33.73 | 9.77 | 0.247 |
|  | Carnivore-piscivore | *Raphiodon vulpinus* | 5 | -31.76 ± 2.21 | 10.47 ± 0.97 | 0.753 ± 0.06 |
|  | Carnivore-piscivore | *Serrasalmus medinai* | 4 | -34.78 ± 4.39 | 9.73 ± 1.5 | 0.196 ± 0.014 |
|  | Carnivore-piscivore | *Serrasalmus spiropleura* | 1 | -23.69 | 11.78 | 0.433 |
|  | Carnivore-piscivore | *Vandellia cirrhosa* | 1 | -25.74 | 14.17 | 0.122 |
|  | Carnivore-piscivore | *Zungaro zungaro* | 1 | -28.8 | 11.93 | 2.763334 |
|  | Invertivore | *Aphyocharax alburnus* | 3 | -26.6 ± 2.37 | 10.14 ± 0.33 | 0.119 ± 0.019 |
|  | Invertivore | *Apteronotus albifrons* | 1 | -28.5 | 8.21 | 0.204 |
|  | Invertivore | *Astyanacinus sp* | 1 | -27.33 | 9.57 | 0.036 |
|  | Invertivore | *Ceratobranchia binghami* | 1 | -24.29 | 9.11 | 0.024 |
|  | Invertivore | *Ceratobranchia elatior* | 2 | -25.9 ± 0.13 | 9.37 ± 0.06 | 0.031 ± 0.027 |
|  | Invertivore | *Characidium etheostoma* | 1 | -28.91 | 9.62 | 0.158 |
|  | Invertivore | *Charax sp* | 1 | -28.33 | 10.54 | 0.463 |
|  | Invertivore | *Cheirodontops sp* | 3 | -30.89 ± 4.43 | 8.89 ± 0.22 | 0.074 ± 0.056 |
|  | Invertivore | *Ctenobrycon hauxwellianus* | 1 | -26.32 | 8.25 | 0.01 |
|  | Invertivore | *Ctenobrycon spilurus* | 1 | -29.57 | 4.74 | 0.09 |
|  | Invertivore | *Eigenmannia trilineata* | 1 | -28.66 | 9.24 | 0.03 |
|  | Invertivore | *Gymnocoriumbus sp* | 1 | -28.12 | 8.25 | 0.048 |
|  | Invertivore | *Jupiaba anteroides* | 4 | -28.42 ± 1.39 | 7.15 ± 0.76 | 0.034 ± 0.008 |
|  | Invertivore | *Knodus moenkhausii* | 1 | -24.99 | 8.89 | 0.036 |
|  | Invertivore | *Knodus sp* | 1 | -25.65 | 8.09 | 0.021 |
|  | Invertivore | *Microcharacidium sp* | 2 | -28.99 ± 1.88 | 7.64 ± 0.77 | 0.047 ± 0.022 |
|  | Invertivore | *Moenkhausia collettii* | 1 | -27.26 | 9.04 | 0.03 |
|  | Invertivore | *Moenkhausia cotinho* | 1 | -25.8 | 10.53 | 0.056 |
|  | Invertivore | *Moenkhausia gr. lepidura* | 1 | -25.08 | 9.48 | 0.114 |
|  | Invertivore | *Moenkhausia oligolepis* | 2 | -27.1 ± 0.25 | 10.06 ± 0.17 | 0.07 ± 0.007 |
|  | Invertivore | *Moenkhausia sp* | 1 | -26.3 | 8.3 | 0.019 |
|  | Invertivore | *Moenkhausia sp 2* | 1 | -25.74 | 8.4 | 0.029 |
|  | Invertivore | *Tatia intermedia* | 1 | -28.71 | 9.62 | 0.05 |
|  | Invertivore | *Tetragonopterus argenteus* | 2 | -26.83 ± 0.04 | 8.48 ± 0.06 | 0.106 ± 0.07 |
|  | Invertivore | *Trachelyopterus galeatus* | 1 | -28.29 | 7.95 | 0.049 |
|  | Omnivore | *Astyanax fasciatus* | 1 | -28.35 | 5.4 | 0.065 |
|  | Omnivore | *Astyanax sp* | 1 | -25.31 | 9.45 | 0.064 |
|  | Omnivore | *Pimelodella aff. cristata* | 5 | -27.35 ± 1.84 | 9.8 ± 0.44 | 0.163 ± 0.06 |
|  | Omnivore | *Pimelodella sp 1* | 4 | -26.58 ± 1.22 | 10.43 ± 0.71 | 0.169 ± 0.129 |
|  | Omnivore | *Pimelodus blochii* | 4 | -27.65 ± 0.68 | 8.88 ± 0.83 | 0.15 ± 0.073 |
|  | Omnivore | *Platydoras costatus* | 1 | -26.65 | 8.67 | 0.076 |
|  | Primary Consumer | *Aphanotorulus unicolor* | 3 | -27.51 ± 2.29 | 7.4 ± 0.11 | 0.053 ± 0.028 |
|  | Primary Consumer | *Liposarcus sp* | 1 | -29.98 | 7.02 | 0.016 |
|  | Primary Consumer | *Odontostilbe fugitiva* | 1 | -29.88 | 8.69 | 0.072 |
|  | Primary Consumer | *Panaque nigrolineatus* | 1 | -28.65 | 8.29 | 0.008 |
|  | Primary Consumer | *Parodon pongoensis* | 1 | -30.12 | 8.75 | 0.036 |
|  | Primary Consumer | *Piaractus brachypomus* | 1 | -22.57 | 6.2 | 0.039 |
|  | Primary Consumer | *Potmorhina latior* | 1 | -35.29 | 9.97 | 0.086 |
|  | Primary Consumer | *Prochilodus rubrotaeniatus* | 12 | -32.82 ± 3.34 | 7.57 ± 0.62 | 0.074 ± 0.03 |
|  | Primary Consumer | *Serrapinnus sp* | 2 | -30.37 ± 1.77 | 9.97 ± 0.83 | 0.089 ± 0.025 |
|  | Primary Consumer | *Steindachnerina argentea* | 1 | -37.97 | 9.34 | 0.016 |
|  | Primary Consumer | *Steindachnerina guentheri* | 2 | -32.12 ± 1.32 | 7.92 ± 0.01 | 0.032 ± 0.011 |
|  | Primary Consumer | *Steindachnerina sp* | 1 | -37.54 | 8.64 | 0.096 |

**Table S2.** Mean (± SD) stable isotope values (δ¹³C and δ¹⁵N) and total mercury concentrations (THg) for basal resources, fish trophic guilds, and bird trophic guilds included in the food web analysis. Sample size (n) indicates the number of individual samples analyzed within each group. Stable isotope ratios are expressed in ‰ relative to international standards (Vienna Pee Dee Belemnite for carbon and atmospheric N₂ for nitrogen). Total mercury (THg) concentrations are reported in mg kg⁻¹ dry weight. Basal resources were not analyzed for mercury and are therefore indicated as unavailable.

| Group | Trophic_guild | mean δ13C | sd δ13C | mean δ15N | sd δ15N | mean THg | sd THg | n |
| --- | --- | --- | --- | --- | --- | --- | --- | --- |
| Allochthonous insects | - | -25.421 | 1.736 | 4.657 | 2.396 | - | - | 53 |
| Allochthonous plants | - | -28.323 | 3.295 | 5.780 | 2.771 | - | - | 24 |
| Aquatic macroinvertebrates | - | -30.398 | 5.287 | 5.396 | 2.219 | - | - | 37 |
| Birds | Aquatic predator | -35.064 | 0.764 | 10.926 | 0.116 | 2.333 | 0.049 | 2 |
|  | FrugiGranivore | -26.642 | 0.836 | 7.448 | 1.364 | 1.351 | 1.994 | 4 |
|  | Invertivore | -26.558 | 0.738 | 8.515 | 0.801 | 1.595 | 1.266 | 8 |
|  | Nectarivore | -27.129 | 1.330 | 9.599 | 0.910 | 6.079 | 7.656 | 8 |
|  | Omnivore | -26.039 | 0.790 | 7.675 | 0.290 | 0.716 | 0.343 | 3 |
| Detritus | - | -27.157 | 2.874 | 5.332 | 2.870 | - | - | 80 |
| Fish | Carnivore-piscivore | -31.000 | 3.851 | 10.371 | 1.512 | 0.382 | 0.253 | 28 |
|  | Invertivore | -27.546 | 2.122 | 8.740 | 1.236 | 0.075 | 0.082 | 36 |
|  | Omnivore | -27.125 | 1.342 | 9.361 | 1.340 | 0.143 | 0.082 | 16 |
|  | Primary Consumer | -31.604 | 3.907 | 7.990 | 1.026 | 0.061 | 0.032 | 27 |
| Macrophytes | - | -27.692 | 6.317 | 4.118 | 2.643 | - | - | 36 |
| Periphyton | - | -30.18 | 3.371 | 14.271 | 5.690 | - | - | 16 |
| Phytoplankton | - | -34.610 | 1.992 | 2.756 | 0.588 | - | - | 3 |

**Detailed Bird trophic guild assignment**

For each bird species, dietary information was compiled from standardized global and regional ornithological sources, including species-level trait databases and natural-history accounts. We first used EltonTraits 1.0 to obtain species-level information on diet composition, foraging stratum, activity period, and body size, which together describe major components of the Eltonian niche of birds (Wilman et al., 2014). This information was cross-checked with species accounts from Birds of the World and, when available, regional Neotropical literature. Guild names and decision rules followed standardized avian diet terminology, particularly the classification scheme proposed by Lopes et al. (2016), which emphasizes the dominant food category and allows mixed categories when species regularly consume more than one major resource type. The use of trophic and trophic-behavioral guilds is also consistent with classical and contemporary studies of Amazonian bird communities, where functional guilds are commonly used to summarize community structure in species-rich assemblages with many rare species and limited direct dietary observations (Terborgh et al., 1990).

Species were assigned to one of five trophic guilds: invertivores, omnivores, frugi-granivores, nectarivores, and aquatic predators. Invertivores included species whose diets are dominated by terrestrial or aerial arthropods, with potential incorporation of aquatic or emergent macroinvertebrates in riparian habitats. Omnivores included species with mixed diets combining animal prey and plant material, without a single clearly dominant resource. Frugi-granivores included species primarily consuming fruits, seeds, or other plant material, while allowing secondary ingestion of arthropods. Nectarivores included hummingbirds and other nectar-feeding species, recognizing that nectar is the dominant energetic resource but that arthropods can represent an important protein source. Aquatic predators included species feeding directly or indirectly on aquatic prey, including fishes and aquatic macroinvertebrates, and species whose foraging ecology is strongly associated with riverine, floodplain, or wetland habitats.

Final assignments were made at the species level following a hierarchical decision rule. First, species were assigned according to the dominant dietary category reported in standardized sources. Second, when species had mixed diets, secondary food categories and foraging habitat were used to determine whether they should be treated as omnivores, frugi-granivores, invertivores, nectarivores, or aquatic predators. Third, isotopic values (δ¹³C and δ¹⁵N) were used only as an ecological consistency check, not as a primary criterion for guild assignment. This validation step allowed us to verify whether species assigned to higher-trophic or aquatic-associated guilds occupied isotopic positions consistent with their expected resource use. Mercury concentrations were not used to assign bird trophic guilds. All species-level assignments, dietary evidence, supporting references, and confidence levels were recorded in a supplementary table to ensure transparency and reproducibility.

**Table S3.** Species-level criteria used for bird trophic guild assignment.

| **Species** | **Dominant diet criterion** | **Secondary diet/resources** | **Aquatic/riparian association criterion** | **Decision rationale** | **Isotopic consistency check / expected pattern** | **Confidence level** | **Assigned guild in final models** |
| --- | --- | --- | --- | --- | --- | --- | --- |
| *Amazilia versicolor* | Nectar | Small arthropods/insects | No direct aquatic dependence; forest/edge nectar resources | Trochilid nectar-feeder; arthropods used as protein source; retained as nectarivore. | Low-to-intermediate δ15N relative to animal predators; δ13C linked to terrestrial/forest plant pathways. | High | Nectarivore |
| *Florisuga mellivora* | Nectar | Small arthropods/insects | No direct aquatic dependence; canopy/edge nectar resources | Hummingbird; nectar dominant with arthropod supplementation. | Low-to-intermediate δ15N; δ13C linked to terrestrial plant pathways. | High | Nectarivore |
| *Glaucis hirsutus* | Nectar | Small arthropods/insects | Often associated with humid/riparian forest edges, but not an aquatic predator | Hermit-like hummingbird; nectar dominant with arthropod supplementation. | Low-to-intermediate δ15N; δ13C linked to terrestrial/riparian plant pathways. | High | Nectarivore |
| *Phaethornis atrimentalis* | Nectar | Small arthropods/insects | No direct aquatic dependence; forest understory nectar resources | Hermit hummingbird; nectar dominant, arthropods secondary protein source. | Low-to-intermediate δ15N; δ13C linked to terrestrial/forest plant pathways. | High | Nectarivore |
| *Phaethornis bourcieri* | Nectar | Small arthropods/insects | No direct aquatic dependence; forest understory nectar resources | Hermit hummingbird; nectar dominant, arthropods secondary protein source. | Low-to-intermediate δ15N; δ13C linked to terrestrial/forest plant pathways. | High | Nectarivore |
| *Phaethornis hispidus* | Nectar | Small arthropods/insects | No direct aquatic dependence; forest/riparian understory nectar resources | Hermit hummingbird; nectar dominant, arthropods secondary protein source. | Low-to-intermediate δ15N; δ13C linked to terrestrial/riparian plant pathways. | High | Nectarivore |
| *Leptotila rufaxilla* | Seeds and fallen fruits | Occasional small invertebrates/plant material | No direct aquatic dependence; ground-foraging in forest and edges | Dove with seed/fruit-dominated ground diet. Current 'Granivore' is compatible with Frugi-granivore. | Low δ15N; δ13C consistent with terrestrial plant/seed-fruit pathway. | High | Frugi-granivore |
| *Chloroceryle aenea* | Small fish | Aquatic insects/crustaceans | Strong; riverine, stream and floodplain-edge foraging | Kingfisher feeding on aquatic prey; direct link to aquatic food web. | High δ15N among birds; δ13C may track aquatic/benthic or fish-supported sources. | High | Aquatic predator |
| *Monasa nigrifrons* | Large insects/arthropods | Occasional small vertebrates | Riparian/forest-edge association possible, but diet is mainly animal prey, not plant omnivory | Nunbird diet is dominated by large arthropods and other animal prey | Intermediate-to-high δ15N relative to other insectivores; δ13C linked to terrestrial/riparian prey. | High | Invertivore |
| *Dendrexetastes rufigula* | Arthropods/insects | Other small invertebrates | No direct aquatic dependence; forest trunk-foraging | Woodcreeper; arthropod-dominated bark/trunk foraging. | Intermediate δ15N; δ13C consistent with forest arthropod pathway. | High | Invertivore |
| *Dendrocolaptes picumnus* | Arthropods/insects | Other small invertebrates; occasional small vertebrates | No direct aquatic dependence; forest trunk-foraging | Woodcreeper with arthropod-dominated foraging. | Intermediate δ15N; δ13C consistent with forest arthropod pathway. | High | Invertivore |
| *Dendrocyncla fuliginosa* | Arthropods/insects | Other small invertebrates | No direct aquatic dependence; forest understory/trunk-foraging | Woodcreeper; arthropod-dominated diet, often associated with ant-following or forest foraging. | Intermediate δ15N; δ13C consistent with forest arthropod pathway. | High | Invertivore |
| *Xiphorhynchus guttatus* | Arthropods/insects | Other small invertebrates; occasional small vertebrates | No direct aquatic dependence; forest trunk-foraging | Woodcreeper diet dominated by arthropods gleaned from bark/trunks. | Intermediate δ15N; δ13C consistent with forest arthropod pathway. | High | Invertivore |
| *Cacicus cela* | Fruit and arthropods | Nectar and other opportunistic resources | No direct aquatic dependence; riparian/edge colonies possible | Icterid with broad mixed diet including fruits and insects; omnivore is appropriate. | Intermediate δ15N; mixed δ13C if fruit and insects contribute substantially. | High | Omnivore |
| *Pipra filicauda* | Small fruits | Arthropods | No direct aquatic dependence; forest understory/subcanopy | Predominantly small-fruit consumers with insects as secondary items | Low-to-intermediate δ15N; δ13C consistent with forest fruit pathway. | High | Frugi-granivore |
| *Thraupis episcopus* | Fruit/plant material | Arthropods; occasional nectar | No direct aquatic dependence; common in edges, forest mosaics and open habitats | Tanager with fruit-dominated mixed diet; insects secondary. Current 'Frugivore' is compatible with Frugi-granivore. | Low-to-intermediate δ15N; δ13C consistent with terrestrial plant/frugivory pathway. | High | Frugi-granivore |
| *Elaenia sp.* | Fruit and arthropods | Small insects and berries; species dependent | No direct aquatic dependence; genus-level classification | Tyrant-flycatchers diet is dominated by small arthorpods. Elaenia spp. commonly combine fruit and insects. | Low-to-intermediate δ15N; mixed δ13C if fruit and insects are both important. | Medium | Invertivore |
| *Poecilotriccus latirostris* | Arthropods/insects | Occasional small invertebrates | No direct aquatic dependence; edge/understory foraging | Tyrant-flycatcher diet dominated by small arthropods captured from vegetation/air. | Intermediate δ15N; δ13C consistent with terrestrial/riparian arthropod pathway. | High | Invertivore |
| *Tolmomyias sp.* | Arthropods/insects | Occasional small invertebrates | No direct aquatic dependence; genus-level classification | Tyrant-flycatcher genus with arthropod-dominated foraging. | Intermediate δ15N; δ13C consistent with terrestrial/riparian arthropods. | Medium | Invertivore |
| *Tyrannus melancholicus* | Arthropods/insects | Fruit, especially seasonally/opportunistically | No direct aquatic dependence; open/riparian edges possible | Flycatcher diet dominated by insects captured aerially; fruit can be secondary but does not dominate guild assignment. | Intermediate δ15N; δ13C consistent with terrestrial/riparian arthropods. | High | Invertivore |
